# Whole-Genome Sequencing and Comparative Genomic Analysis of Potential Biotechnological Strains of Trichoderma harzianum, Trichoderma atroviride*, and* Trichoderma reesei

**DOI:** 10.1101/2022.02.11.479986

**Authors:** Rafaela Rossi Rosolen, Maria Augusta Crivelente Horta, Paulo Henrique Campiteli de Azevedo, Carla Cristina da Silva, Danilo Augusto Sforca, Gustavo Henrique Goldman, Anete Pereira de Souza

## Abstract

*Trichoderma atroviride* and *Trichoderma harzianum* are widely used as commercial biocontrol agents against plant diseases. Recently, *T. harzianum* IOC-3844 (Th3844) and *T. harzianum* CBMAI-0179 (Th0179) demonstrated great potential in the enzymatic conversion of lignocellulose into fermentable sugars. Herein, we performed whole-genome sequencing and assembly of the Th3844 and Th0179 strains. To assess the genetic diversity within the genus *Trichoderma*, the results of both strains were compared with strains of *T. atroviride* CBMAI-00020 (Ta0020) and *T. reesei* CBMAI-0711 (Tr0711). The sequencing coverage value of all genomes evaluated in this study was higher than that of previously reported genomes for the same species of *Trichoderma*. The resulting assembly revealed total lengths of 40 Mb (Th3844), 39 Mb (Th0179), 36 Mb (Ta0020), and 32 Mb (Tr0711). A genome-wide phylogenetic analysis provided details on the relationships of the newly sequenced species with other *Trichoderma* species. Structural variants revealed genomic rearrangements among Th3844, Th0179, Ta0020, and Tr0711 relative to the *T. reesei* QM6a reference genome and showed the functional effects of such variants. In conclusion, the findings presented herein allow the visualization of genetic diversity in the evaluated strains and offer opportunities to explore such fungal genomes in future biotechnological and industrial applications.

## 1 Introduction

Fungi of the genus *Trichoderma* are characterized by their considerable nutritional versatility (Sharma et al., 2019), which allows them to be employed in a wide range of biotechnological applications (Kidwai and Nehra, 2017). For example, *Trichoderma reesei* is the primary fungal source of the industrial cellulases and hemicellulases present in enzymatic cocktails (Bischof et al., 2016). In addition to enzymatic activities, the biocontrol capacity against plant pathogenic fungi has been widely explored in *Trichoderma harzianum* and *Trichoderma atroviride* (Medeiros et al., 2017; Saravanakumar et al., 2017). Recently, *T. harzianum* strains were explored for their enzymatic potential and were demonstrated to be useful for improving lignocellulosic conversion into sugars during second-generation ethanol (2G ethanol) production (Almeida et al., 2021; Delabona et al., 2020; Motta et al., 2021; Zhang et al., 2020).

The phenotypic and ecological heterogeneity across fungi of the same genera is reflected, in part, by the diversity observed within their genomes (Priest et al., 2020). In this way, the diverse and important roles of fungi, as well as the technological advances in next-generation sequencing, have motivated broad efforts to sequence several fungal genomes (Hagestad et al., 2021; Kumar et al., 2021; Nagel et al., 2021; Varga et al., 2019; Wu et al., 2018). Since the genome of *T. reesei* QM6a was first presented (Martinez et al., 2008), *Trichoderma* sequencing studies have increased with the goal of better understanding the biological and ecological roles of *Trichoderma* to improve their applications (Druzhinina et al., 2018; Horta et al., 2014; Kubicek et al., 2011, 2019; Li et al., 2017; Schmoll et al., 2016). Such a growing number of sequenced species may help reveal the molecular basis for the specific features of diverse *Trichoderma* strains.

Previously, the transcriptional profiles of two *T. harzianum* strains, *T. harzianum* IOC-3844 (Th3844) and *T. harzianum* CBMAI-0179 (Th0179), were analyzed under cellulose degradation conditions and compared with those from *T. atroviride* CBMAI-0020 (Ta0020) and *T. reesei* CBMAI-0711 (Tr0711) (Almeida et al., 2021; Horta et al., 2018). Such studies have suggested the great potential of both *T. harzianum* strains as hydrolytic enzyme producers, and this was similar to Tr0711, while Ta0020 showed a low cellulolytic ability. Furthermore, differences in the transcriptional regulation of hydrolytic enzyme expression were observed between Th3844 and Th0179 (Rosolen et al., 2022), highlighting the genetic differences between these strains. Although previous studies investigated the Th3844 genomic regions, which are related to biomass degradation, through bacterial artificial chromosome (BAC) library construction (Crucello et al., 2015; Filho et al., 2017), genomic information regarding the hydrolytic strains of *T. harzianum*, Th3844 and Th0179, remains unclear.

In this study, Pacific Biosciences (PacBio) (Ardui et al., 2018) technology was used to obtain highly contiguous de novo assemblies of Th3844 and Th0179 and identify the genetic variation between them. To expand knowledge on the genetic diversity within the genus *Trichoderma*, the results obtained for *T. harzianum* strains were compared to those from *T. atroviride* and *T. reesei*. The chosen species are appropriate for the study’s goal because while *T. atroviride* is a biocontrol species that is distantly related to the lignocellulolytic species *T. reesei* (Druzhinina et al., 2006), representing a well-defined phylogenetic species (Dodd et al., 2003), *T. harzianum sensu lato* is also commonly used in biocontrol but constitutes a complex of several cryptic species (Chaverri et al., 2015; Druzhinina et al., 2010). Furthermore, we compared the new genomes with publicly available strains, showing the main differences regarding genome features and the superior coverage of the new genomes described here. For phylogenetic comparison, in conjunction with the novel data of the evaluated strains, we used a dataset previously described by Kubicek et al. (Kubicek et al., 2019a).

After performing whole-genome annotation, we investigated the contents of carbohydrate-active enzymes (CAZymes), including glycoside hydrolases (GHs), carbohydrate esterases (CEs), glycosyltransferases (GTs), polysaccharide lyases (PLs), auxiliary activities (AAs), and carbohydrate-binding modules (CBMs) (Cantarel et al., 2009), as well as, based on previous transcriptomes (Almeida et al., 2021; Horta et al., 2018), their gene expression levels under cellulose and glucose growth conditions. We also inspected the secondary metabolite biosynthetic gene clusters (SMGCs) that were distributed among the studied genomes. Biosynthetic gene clusters (BGCs) can be categorized into several classes, including the following: fungal RiPP with POP or UstH peptidase types and a modification (fungal-RiPP); nonribosomal peptide synthetases (NRPSs); NRPS-like fragment (NRPS-like); type I polyketide synthase (T1PKS); terpene; hybrid NRPS, T1PKS; NRPS, T1PKS, beta-lactone; NRPS, T1PKS, NRPS-like; NRPS-like; and terpene, NRPS-like (Brakhage, 2013).

To thoroughly investigate the genetic variability across the four evaluated strains, we explored the structural variants (SVs), which represent a major form of genetic and phenotypic variation that is inherited and polymorphic in species (Mills et al., 2011), between them and *T. reesei* QM6a, the reference genome (Martinez et al., 2008). In addition, by performing a comparative genomic analysis across the genus *Trichoderma* and more evolutionarily distant genera, the orthologs and the orthogroups across them were identified, and the rooted gene tree based on the single-copy orthologs was inferred.

The genomic resources we provide herein significantly extend our knowledge regarding the evolution and basic biology of the evaluated strains, which may increase their biotechnological employment. The results from this study might also increase the availability of genomic data, which can be used for comparative studies aiming to correlate the phenotypic differences in the genetic diversity of *Trichoderma* species; therefore, the study may help improve the search for enzymes with enhanced properties and provide aid toward improving the production of chemicals and enzymes in such fungi.

## 2 Materials and methods

### 2.1 Fungal strains and culture conditions

The species originated from the Brazilian Collection of Environment and Industry Microorganisms (CBMAI), which is located in the Pluridisciplinary Center for Chemical, Biological, and Agricultural Research (CPQBA) at the University of Campinas (UNICAMP), Brazil. The identity of *Trichoderma* isolates was authenticated by CBMAI based on phylogenetic studies of their internal transcribed spacer (ITS) region, translational elongation factor 1 (*tef1*), and RNA polymerase II (*rpb2*) marker gene. The phylogenetic tree based on the ITS region was previously published (Rosolen et al., 2022); moreover, the methodology used to model the phylogenetic trees based on the *tef1* and *rpb2* marker genes is described in Supplementary Material 1. Briefly, Th3844, Th0179, Ta0020, and Tr0711 strains were cultivated on potato dextrose agar (PDA) solid medium (ampicillin 100 µg/ml and chloramphenicol 34 µg/ml) for 3 days at 28 °C. Conidia were harvested, and an initial spore solution was used to inoculate 500 mL of potato dextrose broth (PDB) medium. The cultivation process was performed in biological triplicates for 72□h at 28 °C and 200 rpm for all evaluated strains. Then, mycelial samples were harvested using Miracloth (Millipore), frozen using liquid nitrogen, and stored at −80 °C. Frozen material was used for DNA extraction.

### 2.2 DNA extraction and sequencing

The ground fungal tissue was suspended in 800 μl of lysis buffer (1 M Tris-HCl [pH 8.0], 0.5 M EDTA [pH 8.0], 5 M NaCl, N-lauroylsarcosine, β-mercaptoethanol, and H_2_O), and phenol:chloroform:isoamyl alcohol (25:24:1) (Sigma, US) was then added. After centrifugation at 4□°C and 15,871 x g for 10□min, the aqueous layer was collected, and genomic DNA was precipitated through the addition of isopropanol. DNA was harvested by centrifugation at 4□°C and 15,871 x g for 10□min, and the pellet was washed with 70% ethanol and centrifuged at 4□°C and 15,871 x g for 5 min. After a second wash with 95% ethanol and centrifugation at 4□°C and 15,871 x g for 5 min, the pellet was dried at room temperature and dissolved in TE buffer. Before the quality control steps, the DNA was subjected to RNase treatment, which involved the addition of 2 µl of RNase to the TE buffer (final concentration of 50 µg/mL) and incubation of the samples for 1 h at room temperature before storage. The methodology used for DNA extraction was based on previously established protocols (Oliveira et al., 2015; Peterson et al., 2000) and adapted according to our fungal samples.

The quantity of the extracted gDNA was determined by measuring the absorbance at 260 nm using a NanoDrop 1000 spectrophotometer (Thermo Fisher Scientific) and Qubit Fluorometer (Thermo Fisher Scientific). The quality of extracted gDNA was assessed through 0.8% agarose gel electrophoresis. HiFi sequencing libraries were prepared according to the PacBio protocol (Wenger et al., 2019), and sequencing was performed at the Arizona Genomics Institute (AGI; Tucson, USA) using a SMRT DNA sequencing system from PacBio (PacBio RSII platform).

### 2.3 Genome assembly

The data were transferred to a local server, and the genomes were assembled de novo using Canu software (v.2.1) (-pacbio – hifi, and a genome estimate equal to 40 Mb for all evaluated strains), which was developed for long-read sequencing (Koren et al., 2017). Genome integrity was assessed using the Quality Assessment Tool (QUAST) (Gurevich et al., 2013) (v.5.0.2) and Benchmarking Universal Single-Copy Orthologs (BUSCO) (Simao et al., 2015) (v.4.1.4) tools. The Nucmer alignment tool from the MUMmer (v.4.0.0beta2) toolbox (Kurtz et al., 2004; Marcais et al., 2018) was used to perform the whole-genome alignments between the evaluated strains.

### 2.4 Gene prediction and functional annotation

Gene prediction was performed using AUGUSTUS (v.3.3.3) (Stanke et al., 2006) through gene models, which were built from *T. harzianum* T6776, *T. atroviride* IMI206040, and *T. reesei* QM6a (TrainAugustus (v.3.3.3)), together with MAKER (Cantarel et al., 2008) (v.2.31.11). Such programs are implemented on the Galaxy platform. The predicted genes were functionally annotated by searching for homologous sequences in the UniProt (The UniProt Consortium, 2021), eggNOG-mapper v.2 (Cantalapiedra et al., 2021), and Protein Annotation with Z score (PANNZER2) (Toronen et al., 2018) databases. Transmembrane proteins were predicted using TMHMM v.2.0 (Krogh et al., 2001). For the annotation of CAZymes, we used CDSs as homology search queries against the database of the dbCAN2 server (Zhang et al., 2018), which integrates the (I) DIAMOND (E-value < 1e-102) (Buchfink et al., 2015), (II) HMMER (E-value < 1e-15, coverage > 0.35) (Finn et al., 2011), and Hotpep (frequency > 2.6, hits > 6) (Busk et al., 2017) tools. We considered all CDSs as true hits if they were predicted by at least two tools. Coverages were estimated with QualiMap (Okonechnikov et al., 2016) (v.2.2.2c) using minimap2 (Li, 2018) v. 2.17 + galaxy4, which were both implemented on the Galaxy platform (Afgan et al., 2018). SMGCs in the Th3844, Th0179, Ta0020, and Tr0711 genomes were predicted using antiSMASH fungal version v.6.1.0 (Blin et al., 2021).

### 2.5 Ortholog identification and clustering

The proteomes of Th3844, Th0179, Ta0020, and Tr0711 were compared with those of 18 *Trichoderma* spp. proteomes that are available in NCBI databases. *Fusarium* spp., *Aspergillus* spp., and *Neurospora* spp. were used as outgroups. For this analysis, we used the software OrthoFinder (Emms and Kelly, 2015, 2019) v2.5.2, which clustered the protein sequences of fungi into orthologous groups and allowed the phylogenetic relationships between them to be identified. The consensus species tree was inferred using the STAG algorithm (Emms and Kelly, 2018) and rooted using the STRIDE algorithm (Emms and Kelly, 2017), which was implemented in the OrthoFinder program. The resulting tree from the OrthoFinder analysis was visualized and edited using Interactive Tree of Life (iTOL) v6 (Letunic and Bork, 2007).

### 2.6 Long-read structural variant analysis

SVs were identified by aligning the PacBio HiFi reads from Th3844, Th0179, Ta0020, and Tr0711 with the *T. reesei* QM6a reference genome (Martinez et al., 2008) using the software Map with BWA-MEM (Li and Durbin, 2010) v.0.7.17.2 with (-x pacbio, sorted by chromosomal coordinates). The duplicate reads in the BAM file were identified and marked using the tool MarkDuplicates (Broad Institute) v.2.18.2.2. Variants were called using Sniffles (Sedlazeck et al., 2018) v.1.0.12+galaxy0, allowing for a minimum support of 10 (--min_support), maximum number of splits of 7 (--max_num_splits), maximum distance of 1000 (--max_distance), minimum length of 30 (-- min_length), minimum mapping quality of 20 (--minmapping_qual), and CCS reads option (--ccs_reads). SVs were annotated using SnpEff (v.4.3+T. galaxy1) (Cingolani et al., 2012), which allowed the effects of variants in genome sequences to be categorized. These tools were implemented using the Galaxy platform (Afgan et al., 2018).

### 2.7 Expression profile of CAZymes

The gene expression profile of the CAZymes identified in the assembled genomes was investigated using data from previous transcriptomes (Almeida et al., 2021; Horta et al., 2018). The gene expression of CAZymes from Th3844, Th0179, Ta0020, and Tr0711 was modeled using transcripts per million (TPM) data, which were validated through qPCR and described by Almeida et al. (2021) and Horta et al. (2018). This gene expression dataset was obtained from three biological replicates under cellulose and glucose growth conditions.

To search for the corresponding homologs, we mapped all transcripts from (I) Th3844 and Th0179, (II) Ta0020, and (III) Tr0711 against the nucleotide sequences from (I) *T. harzianum* T6776 (Baroncelli et al., 2015), (II) *T. atroviride* IMI 206040 (Kubicek et al., 2011), and *T. reesei* v2.0 (Martinez et al., 2008), respectively. We then selected the transcripts corresponding to the CAZymes identified in the assembled genomes, and their expression patterns under cellulose and glucose growth conditions were evaluated. The corresponding orthologs of CAZyme-encoding genes among the genomes of Th3844, Th0179, Ta0020, and Tr0711 were obtained using the results from the OrthoFinder analysis, and the results were visualized using the R package pheatmap (Kolde, 2019).

## 3 Results

### 3.1 Molecular phylogeny of the evaluated Trichoderma spp. isolates, strain cultivation, and evaluation of extracted DNA

For the molecular identification of the fungi used in this study, we previously investigated the phylogenetic relationships among Th3844, Th0179, Ta0020, and Tr0711 based on the ITS sequences (Rosolen et al., 2022). Here, we modeled phylogenetic trees based on the *rpb2* and *tef1* sequences of 16 and 26 *Trichoderma* spp., respectively, including those of our study strains (Supplementary Material 1: Supplementary Fig. 1). According to the results, high genetic proximity was observed between Th3844 and Th0179. In contrast, both strains were phylogenetically distant from Ta0020 and Tr0711, which were grouped with other *T. atroviride* and *T. reesei* strains, respectively. *Neurospora* spp. and *Fusarium* spp. were used as outgroups of the *rpb2* phylogenetic tree. No outgroups were used in the *tef1* phylogenetic tree because the orthologs in the most distant genera, including *Neurospora* spp. and *Fusarium* spp., presented significantly different transcript sequences, which did not allow alignment between their nucleotide sequences with those of *Trichoderma* spp.

First, we cultivated Th3844, Th0179, Ta0020, and Tr0711. After 3 days of culture at 28 °C, Tr0711 presented the highest production of spores, followed by Th0179, Th3844, and Ta0020 (Supplementary Material 1: Supplementary Fig. 2). Additionally, after 7 days of culture at 28 °C, the four strains showed different growth patterns, i.e., Th3844 (Fig. 1A) presented a production of spores similar to Th0179 (Fig. 1B) and Tr0711 (Fig. 1C), and Ta0020 (Fig. 1D) showed slow sporulation. The DNA from the evaluated *Trichoderma* isolates was then extracted, and its integrity and quality were assessed (Supplementary Material 1: Supplementary Table 1 and Supplementary Material 1: Supplementary Fig. 3).

**Fig. 1.**
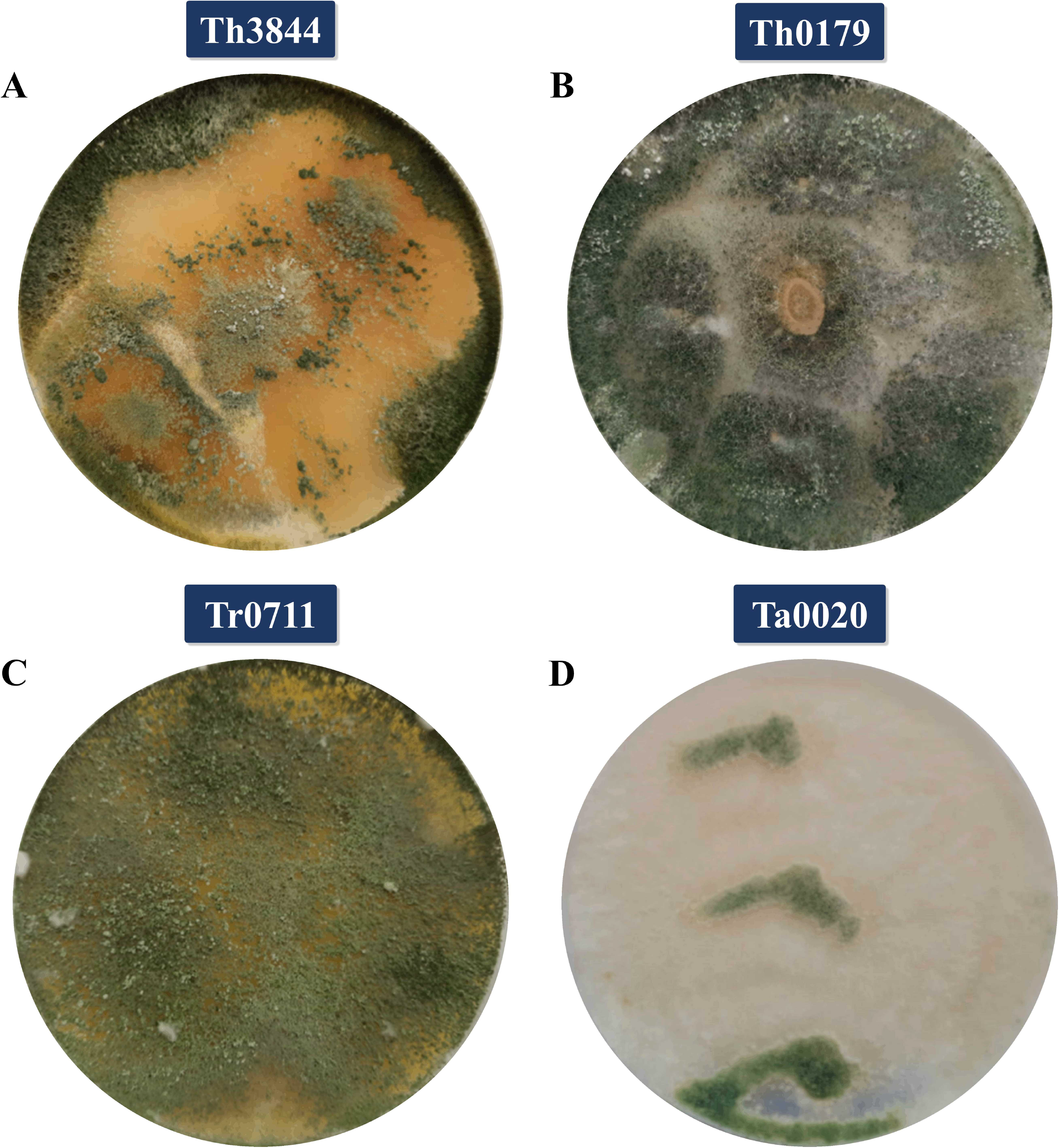
*Trichoderma* isolates evaluated in this study. (a) Th3844, (b) Th0179, (c) Tr0711, and (d) Ta0020 cultivated on potato dextrose broth (PDA) solid medium at 28 °C. The four strains showed a different growth pattern, i.e., Th3844 (a) presented a production of spores similar to Th0179 (b) and Tr0711 (c), and Ta0020 (d) showed slow sporulation. Ta0020: *T. atroviride* CBMAI-0020; Th0179: *T. harzianum* CBMAI-0179; Th3844: *T. harzianum* IOC-3844; Tr0711: *T. reesei* CBMAI-0711

### 3.2 Assembled genomic features and general comparison across *Trichoderma* spp

In the present study, we introduced the whole-genome sequences of Th3844, Th0179, Ta0020, and Tr0711 (Table 1). Overall, the genomes of the evaluated *Trichoderma* spp. varied in the number of contigs (14–26), sizes (32– 40□Mb) and gene contents (8,796–11,322 genes). In comparison with the other strains, Th0711 contains the smallest gene repertoire, while Th3844 contains the highest gene repertoire. To assess the completeness and integrity of the assembled genomes, BUSCO analysis was performed. For all evaluated strains, over 90% of genes were complete. Although the genome of Th3844 presented a considerable number of missing genes (9.7%), its assembled genome exhibited a lower degree of fragmentation compared to that of other genomes.

**Table 1.**
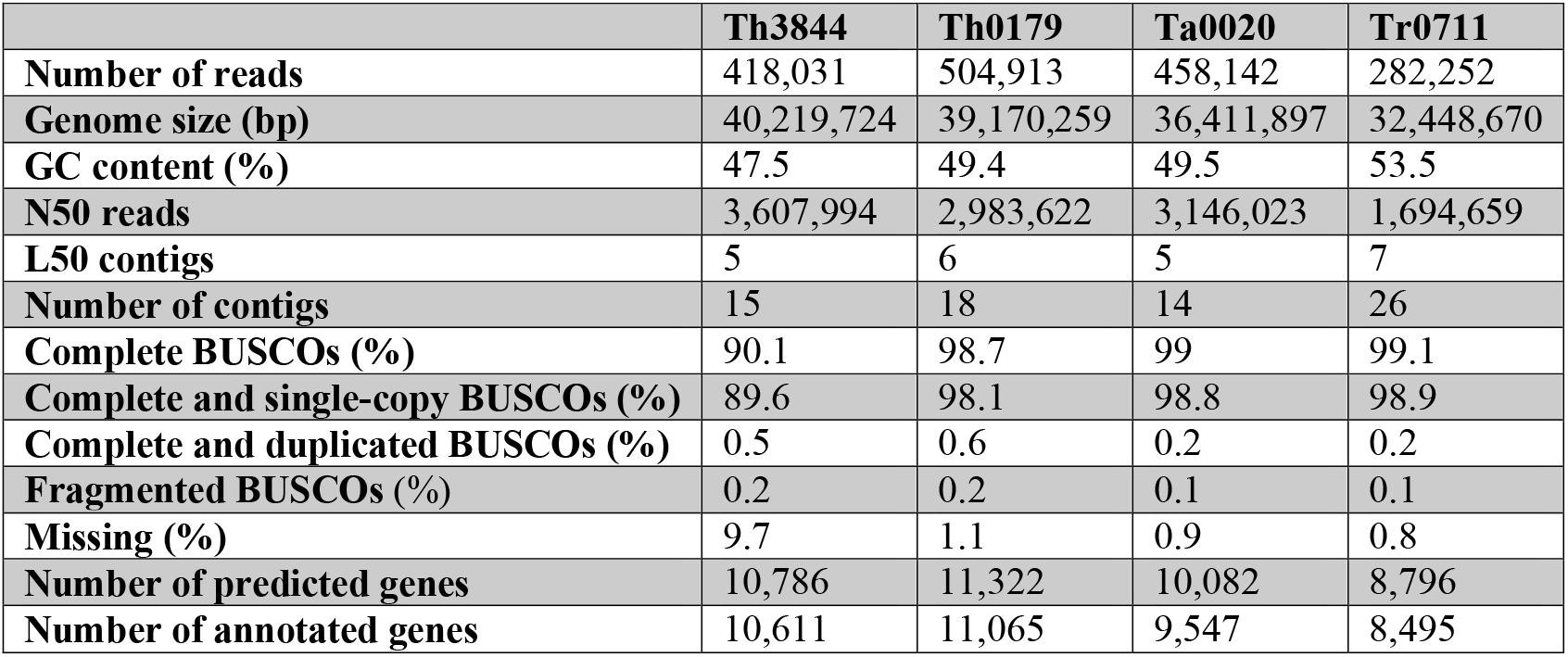
Genome assembly and annotation statistics.

|  | **Th3844** | **Th0179** | **Ta0020** | **Tr0711** |
| --- | --- | --- | --- | --- |
| **Number of reads** | 418,031 | 504,913 | 458,142 | 282,252 |
| **Genome size (bp)** | 40,219,724 | 39,170,259 | 36,411,897 | 32,448,670 |
| **GC content (%)** | 47.5 | 49.4 | 49.5 | 53.5 |
| **N50 reads** | 3,607,994 | 2,983,622 | 3,146,023 | 1,694,659 |
| **L50 contigs** | 5 | 6 | 5 | 7 |
| **Number of contigs** | 15 | 18 | 14 | 26 |
| **Complete BUSCOs (%)** | 90.1 | 98.7 | 99 | 99.1 |
| **Complete and single-copy BUSCOs (%)** | 89.6 | 98.1 | 98.8 | 98.9 |
| **Complete and duplicated BUSCOs (%)** | 0.5 | 0.6 | 0.2 | 0.2 |
| **Fragmented BUSCOs (%)** | 0.2 | 0.2 | 0.1 | 0.1 |
| **Missing (%)** | 9.7 | 1.1 | 0.9 | 0.8 |
| **Number of predicted genes** | 10,786 | 11,322 | 10,082 | 8,796 |
| **Number of annotated genes** | 10,611 | 11,065 | 9,547 | 8,495 |

Considering the genome sequencing coverage, genome size, GC content, length metrics (N50 and L50 values), assembly level, and number of genes, the genomes of Th3844, Th0179, Ta0020, and Tr0711 were compared to other fungal genome references (Baroncelli et al., 2015; Chung et al., 2021; Kubicek et al., 2011; Li et al., 2017; Martinez et al., 2008) (Table 2). All *T. harzianum* genomes were similar in size and GC content. The same profile was observed for the *T. atroviride* and *T. reesei* genomes. Large differences were found for the genome sequencing coverage, in which Th3844, Th0179, Ta0020, and Tr0711 presented higher values than those reported for other strains in the literature (Baroncelli et al., 2015; Kubicek et al., 2011; Li et al., 2017). In regard to quality, except for Tr0711, the genomes assembled in this study showed a lower degree of fragmentation compared to those previously available (Baroncelli et al., 2015; Chung et al., 2021; Kubicek et al., 2011; Li et al., 2017; Martinez et al., 2008).

**Table 2.** Comparison of the genome features of Trichoderma spp. genomes.

| Species | Strain | Cove<br>rage | NCBI accession | Genome<br>size<br>(Mb) | GC<br>content<br>(%) | N50<br>reads | L50<br>contigs | Assembly<br>level | Genes | Reference |
| --- | --- | --- | --- | --- | --- | --- | --- | --- | --- | --- |
| *T. harzianum* | IOC-3844 | 164 x | SAMN23309297 | 40 | 47.5 | 3,607,994 | 5 | Contigs (15) | 10,786 | This study |
| *T. harzianum* | CBMAI-0179 | 219 x | SAMN23309298 | 39 | 49.4 | 2,983,622 | 6 | Contigs (18) | 11,322 | This study |
| *T. harzianum* | T6776 | 85 x | SAMN02851310 | 39 | 48.5 | 68,846 | 178 | Scaffolds<br>(1,572) | 11,501 | (Baroncelli et al., 2015) |
| *T. harzianum* | CBS 226.95 | 120 x | SAMN00761861 | 41 | 47.6 | 2,414,909 | 7 | Scaffolds (532) | 14,269 | - |
| *T. virens* | Gv29–8 | 8 x | SAMN02744059 | 39 | 49.2 | 1,836,662 | 8 | Scaffolds (93) | 12,405 | (Kubicek et al., 2011) |
| *T. simmonsii* | GH-Sj1 | - | SAMN15516371 | 40 | 48.13 | 6,451,197 | 3 | Scaffolds (7) | 13,296 | (Chung et al., 2021) |
| *T. atroviride* | CBMAI-0020 | 229 x | SAMN23309299 | 36 | 49.5 | 3,146,023 | 5 | Contigs (14) | 10,082 | This study |
| *T. atroviride* | IMI206040 | 8 x | SAMN02744066 | 36 | 49.7 | 2,007,903 | 6 | Contigs (29) | 11,809 | (Kubicek et al., 2011) |
| *T. reesei* | CBMAI-0711 | 143 x | SAMN23309300 | 32 | 53.5 | 1,694,659 | 7 | Contigs (26) | 8,796 | This study |
| *T. reesei* | QM6a | 80 x | SAMN05250858 | 35 | 51.0 | 18,236 | 3 | Chromosomes<br>(7) | 10,877 | (Li et al., 2017) |
| *T. reesei* | QM6a | 9 x | SAMN02746107 | 33 | 52.8 | 1,219,543 | 9 | Scaffolds (77) | 9,109 | (Martinez et al., 2008) |

We also performed alignment analyses to evaluate the similarities and variations between the genomes of the studied strains (I) and *T. reesei* QM6a, which is a model organism for the lignocellulose deconstruction system and has a genome that is assembled at the chromosomal level (Li et al., 2017) (Fig. 2), and (II) and their respective reference genomes (Supplementary Material 1: Supplementary Fig. 4). The profile of alignment across the genomes illustrated the degree of divergence across the studied strains with *T. reesei* QM6a and with the closest related strain of each evaluated fungus. For each evaluated strain, we observed alignment in different regions of the *T. reesei* QM6a reference genome. Using *T. reesei* QM6a as a reference genome, we found a total of (I) 12% aligned bases for Th3844, (II) 13% aligned bases for Th0179, (III) 8% aligned bases for Ta0020, and (IV) 95% aligned bases for Tr0711. The total percentage of aligned bases for the Th3844 and Th0179 genomes with the *T. harzianum* T6776 genome was approximately 83.5% and 89%, respectively. In addition, for the Ta0020 genome, the total percentage of aligned bases to the *T. atroviride* IMI206040 reference genome was approximately 88%.

**Fig. 2.**
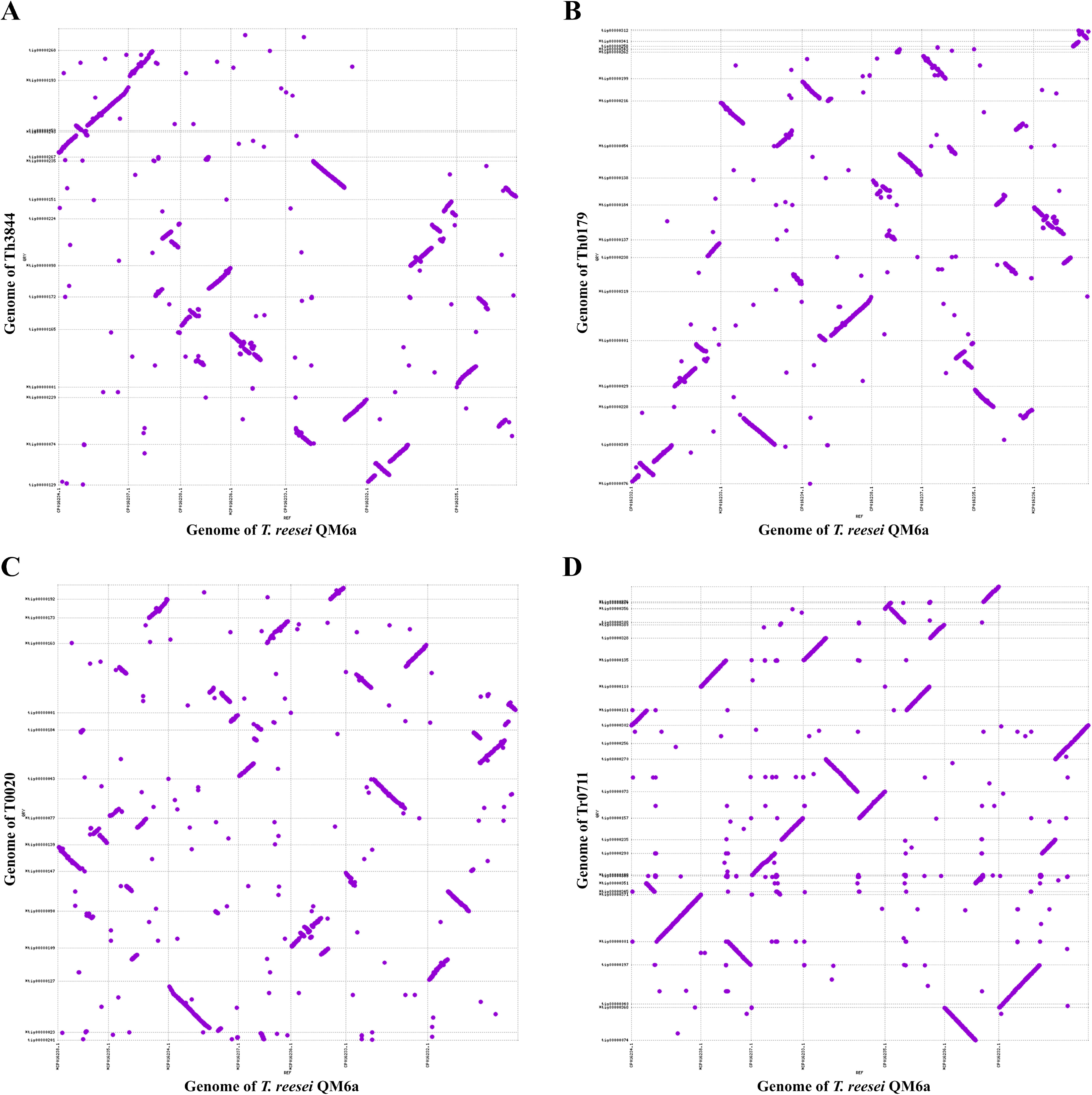
Comparisons between the genomes of the analyzed *Trichoderma* isolates and the *T. reesei* QM6a reference genome. Dot plots of the assemblies of (a) Th3844, (b) Th0179, (c) Ta0020, and (d) Tr0711 that were generated by Canu (y-axis) against those from *T. reesei* QM6a (x-axis) that are available in the NCBI database. Ta0020: *T. atroviride* CBMAI-0020; Th0179: *T. harzianum* CBMAI-0179; Th3844: *T. harzianum* IOC-3844; Tr0711: *T. reesei* CBMAI-0711

### 3.3 Functional annotation, secondary metabolite gene cluster diversity, and CAZyme distribution and quantification in evaluated *Trichoderma* spp. genomes

After performing the genome assembly and gene prediction steps, functional annotation was accomplished by a homology search. The functional category distribution regarding the clusters of orthologous groups of proteins (COGs) (Tatusov et al., 2000) is shown in Fig. 3 and Supplementary Material 1: Supplementary Table 2. Disregarding the (S) function unknown category, the top 5 functional categories were (G) carbohydrate metabolism and transport, (O) posttranslational modification, protein turnover, chaperone functions, (Q) secondary structure, (E) amino acid transport and metabolism, and (U) intracellular trafficking, secretion, and vesicular transport. Overall, the evaluated strains seem to have similar COG assignment profiles. The complete functional annotation of the four genomes is available in Supplementary Material 2: Supplementary Table 3.

**Fig. 3.**
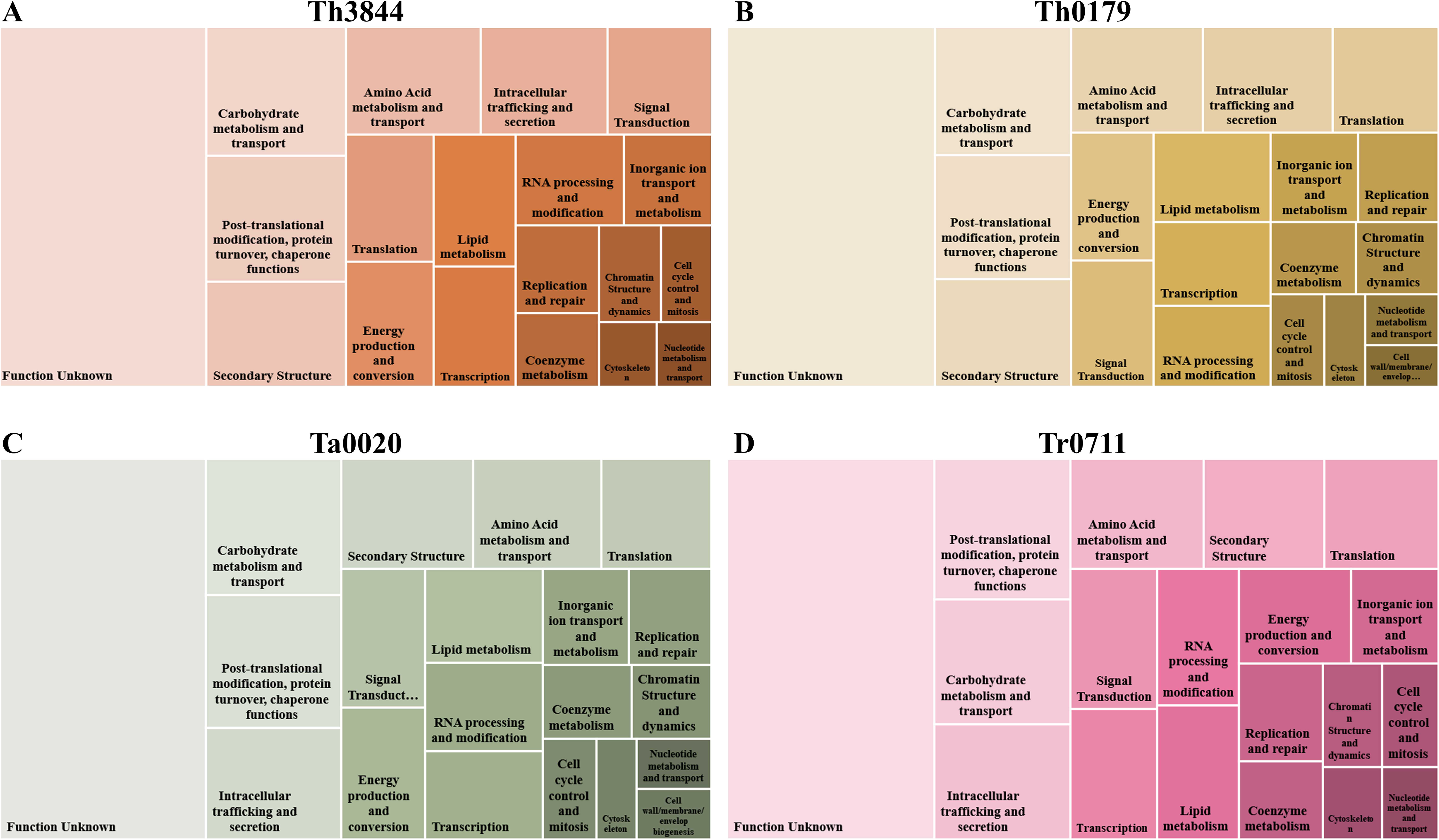
COG functional category distribution of the *Trichoderma* isolates considered. The plot shows the number of genes in the genomes of (a) Th3844, (b) Th0179, (c) Ta0020, and (d) Tr0711, in which a COG classification was obtained. The size of the boxes represents the abundance of the genes at the level of individual COG families. Only the COG functional categories with more than a hundred counts were represented. Ta0020: *T. atroviride* CBMAI-0020; Th0179: *T. harzianum* CBMAI-0179; Th3844: *T. harzianum* IOC-3844; Tr0711: *T. reesei* CBMAI-0711

AntiSMASH predicted the presence of different types of BGCs encoding potential secondary metabolites (SMs). The BGCs of Th3844, Th0179, Ta0020, and Tr0711 are listed in Supplementary Material 3: Supplementary Table 4. In total, 48 (I), 63 (II), 42 (III), and 30 (IV) BGCs were predicted for (I) Th3844, (II) Th0179, (III) Ta0020, and (IV) Tr0711, respectively. Among these, (I) 17, (II) 21, (III) 10, and (IV) 9 clusters showed similarity to BGCs with a known function. For Th3844, five gene clusters containing 100% of the genes from the known cluster were identified (Table 3), whereas for the other strains, six (Th0179), four (Ta0020), and three (Tr0711) gene clusters were recognized.

**Table 3.**
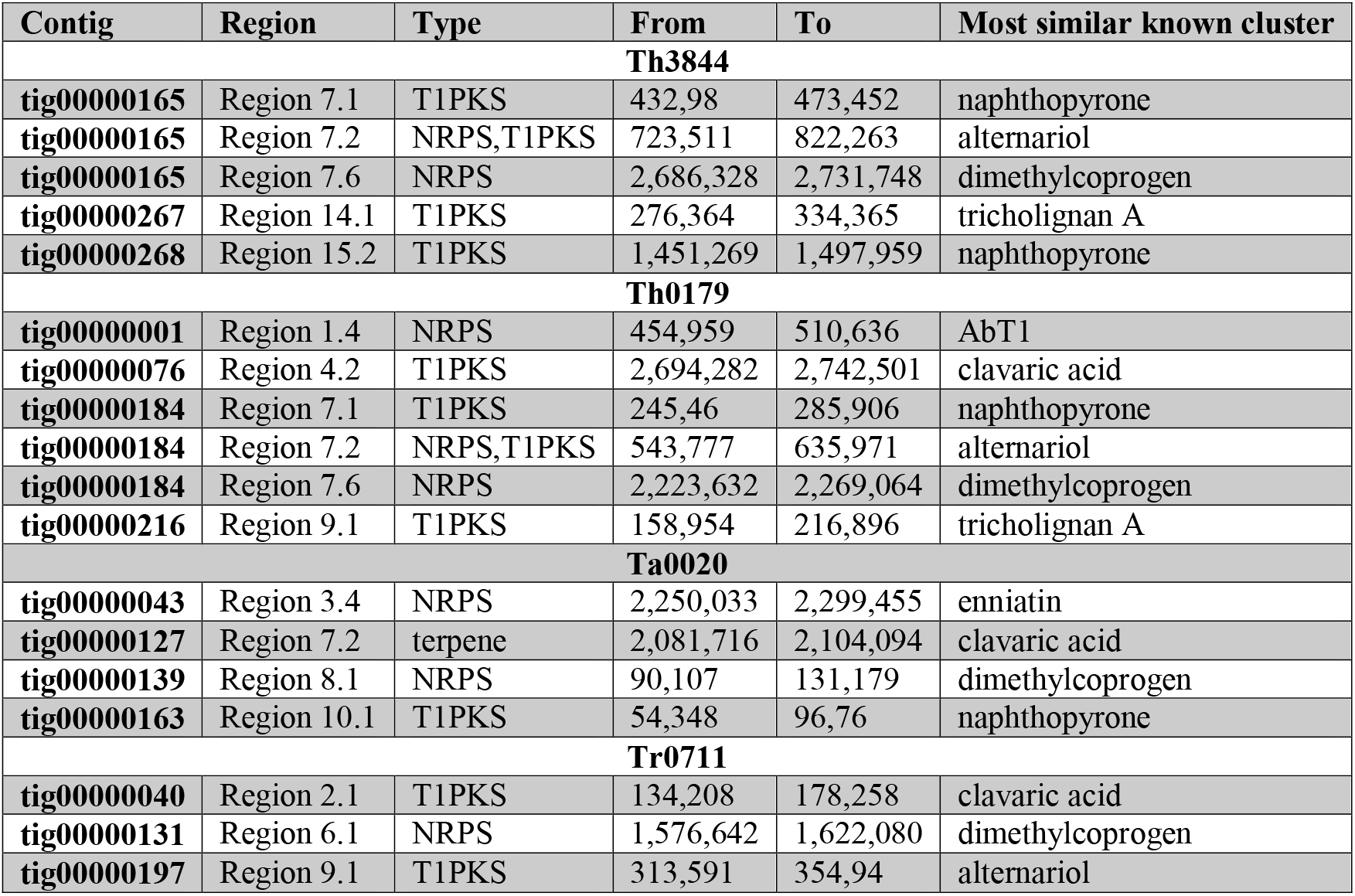
Potential biosynthetic gene clusters in Th3844, Th0179, Ta0020, and Tr0711 showing 100% similarity with the known clusters predicted by antiSMASH.

| Contig | Region | Type | From | To | Most similar known cluster |
| --- | --- | --- | --- | --- | --- |
| **Th3844** |  |  |  |  |  |
| **tig00000165** | Region 7.1 | T1PKS | 432,98 | 473,452 | naphthopyrone |
| **tig00000165** | Region 7.2 | NRPS,T1PKS | 723,511 | 822,263 | alternariol |
| **tig00000165** | Region 7.6 | NRPS | 2,686,328 | 2,731,748 | dimethylcoprogen |
| **tig00000267** | Region 14.1 | T1PKS | 276,364 | 334,365 | tricholignan A |
| **tig00000268** | Region 15.2 | T1PKS | 1,451,269 | 1,497,959 | naphthopyrone |
| **Th0179** |  |  |  |  |  |
| **tig00000001** | Region 1.4 | NRPS | 454,959 | 510,636 | AbT1 |
| **tig00000076** | Region 4.2 | T1PKS | 2,694,282 | 2,742,501 | clavaric acid |
| **tig00000184** | Region 7.1 | T1PKS | 245,46 | 285,906 | naphthopyrone |
| **tig00000184** | Region 7.2 | NRPS,T1PKS | 543,777 | 635,971 | alternariol |
| **tig00000184** | Region 7.6 | NRPS | 2,223,632 | 2,269,064 | dimethylcoprogen |
| **tig00000216** | Region 9.1 | T1PKS | 158,954 | 216,896 | tricholignan A |
| **Ta0020** |  |  |  |  |  |
| **tig00000043** | Region 3.4 | NRPS | 2,250,033 | 2,299,455 | enniatin |
| **tig00000127** | Region 7.2 | terpene | 2,081,716 | 2,104,094 | clavaric acid |
| **tig00000139** | Region 8.1 | NRPS | 90,107 | 131,179 | dimethylcoprogen |
| **tig00000163** | Region 10.1 | T1PKS | 54,348 | 96,76 | naphthopyrone |
| **Tr0711** |  |  |  |  |  |
| **tig00000040** | Region 2.1 | T1PKS | 134,208 | 178,258 | clavaric acid |
| **tig00000131** | Region 6.1 | NRPS | 1,576,642 | 1,622,080 | dimethylcoprogen |
| **tig00000197** | Region 9.1 | T1PKS | 313,591 | 354,94 | alternariol |

Overall, among the tested strains, Th0179 showed the highest number of BGCs, followed by Th3844, Ta0020, and Tr0711 (Fig. 4). Additionally, the T1PKS class was the most represented among all evaluated strains. Interestingly, the NRPS class was more highly represented in Th0179 than in the other strains.

**Fig. 4.**
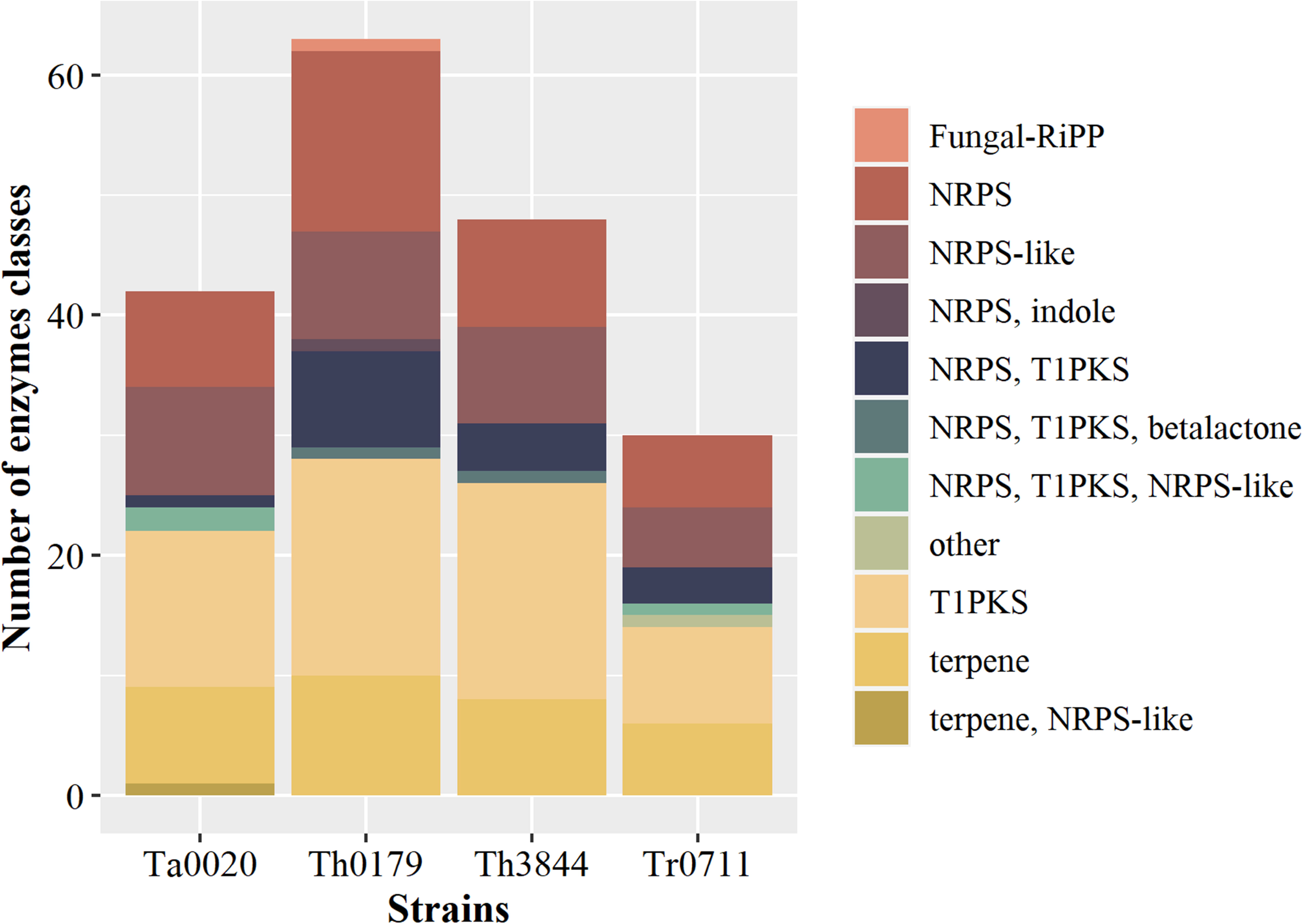
Quantitative comparison of the enzyme classes involved in SM synthesis. Different classes of enzymes related to the production of SMs were predicted from the Th3844, Th0179, Ta0020, and Tr0711 genomes. Among all the strains, Th0179 exhibited the highest number of BGCs, followed by Th3844, Ta0020, and Tr0711. Although the T1PKS class was the most represented among all evaluated strains, the NRPS class was more highly represented in Th0179 than in the other strains. Fungal-RiPP: fungal RiPP with POP or UstH peptidase types and a modification; NRPS: nonribosomal peptide synthetase; T1PKS: type I polyketide synthase; Ta0020: *T. atroviride* CBMAI-0020; Th0179: *T. harzianum* CBMAI-0179; Th3844: *T. harzianum* IOC-3844; Tr0711: *T. reesei* CBMAI-0711

Therefore, we classified the total CAZyme content at the genomic level among the new potential biotechnological strains evaluated in this study (Supplementary Material 4: Supplementary Table 5). To detect similarities and differences between the strains, their CAZyme profiles were compared (Fig. 5A). Overall, the results indicated great variability in the CAZyme content related to the biomass degradation process and mycoparasitism among the Th3844, Th0179, Ta0020, and Tr0711 genomes, as explored next. Among the main CAZyme classes detected in all strains, GHs were overrepresented, and Th3844 (256) and Th0179 (252) had the highest number, followed by Ta0020 (230) and Tr0711 (184). This CAZyme class was followed by GTs as follows: (I) 88 (Th3844), (II) 90 (Th0179), (III) 90 (Ta0020), and (IV) 86 (Tr0711). However, the analysis of the CAZymes with predicted signal peptides revealed that only a few GTs were secreted, as follows: (I) 8 (Th3844), (II) 9 (Th0179), (III) 7 (Ta0020), and (IV) 4 (Tr0711) (Fig. 5B). The distribution of the different CAZyme families among strains was investigated and is available in Supplementary Material 1: Supplementary Fig. 5.

**Fig. 5.**
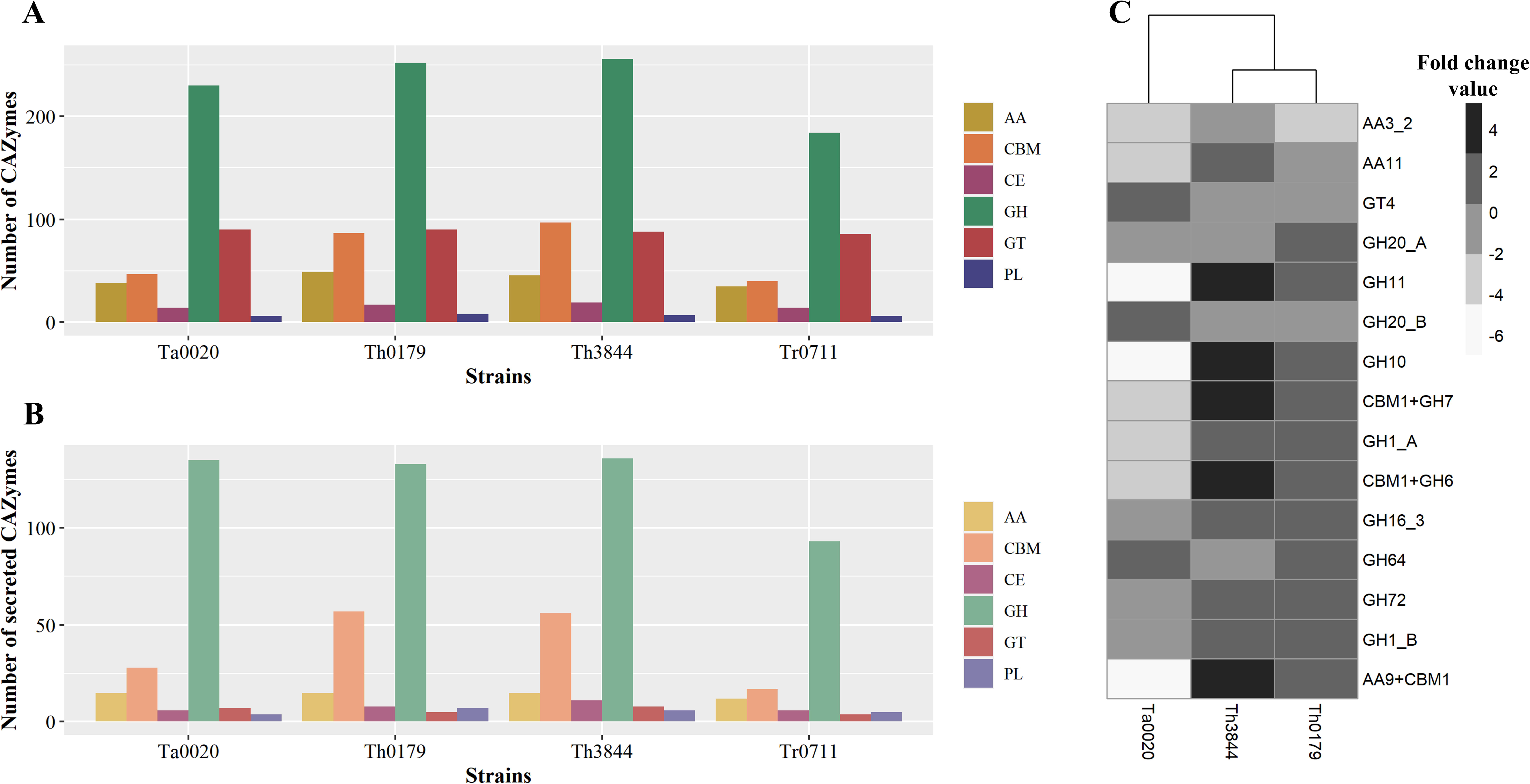
Distribution of CAZymes in *Trichoderma* spp. and evaluation of their expression in Th3844, Th0179, and Ta0020 cells by RNA-Seq. (a) The predicted CAZymes from the assembled genomes were classified according to the CAZy database. (b) The secreted CAZymes were grouped according to their CAZyme class. (c) Heatmap plotted based on the CAZymes of Th3844, Th0179, and Ta0020 showing upregulated and downregulated expression under cellulose and glucose growth conditions, as indicated by an increase or decrease in the gray color in the subtitle. The expression of the CAZyme classes of enzymes with orthologs identified between Th3844, Th0179, and Ta0020 was quantified. CAZymes: carbohydrate-active enzymes; Ta0020: *T. atroviride* CBMAI-0020; Th0179: *T. harzianum* CBMAI-0179; Th3844: *T. harzianum* IOC-3844; Tr0711: *T. reesei* CBMAI-0711. AA: auxiliary activity; CBM: carbohydrate-binding module; EC: carbohydrate esterase; GH: glycosyl hydrolase; GT: glycosyl transferase; PL: polysaccharide lyase

We also investigated the expression profile of the CAZymes identified in the assembled genomes under cellulose and glucose growth conditions. Considering the differentially expressed genes (DEGs) under cellulose growth conditions, we found 35, 78, 31, and 14 CAZyme genes for Th0179, Th3844, Ta0020, and Tr0711, respectively. The classification of the CAZyme genes of Th0179, Th3844, Ta0020, and Tr0711 under cellulose fermentative conditions along with their gene ID number, protein product number, corresponding homolog in the reference genome, fold change values, e-values, and potentially secreted protein are described in Supplementary Material 1: Supplementary Tables 6-9, respectively. The DEGs under cellulose growth conditions that presented orthologs across Th3844, Th0179, and Ta0020 are represented in Fig. 5C, and the identification of the corresponding ortholog for each strain is shown in Supplementary Material 1: Supplementary Table 10. Curiously, Tr0711 showed only one CAZyme-encoding gene from the AA3_2 subfamily with orthologs in the other evaluated strains (Th3844_011000, Th0179_011191, Ta0020_000129, and Tr0711_006613). Overall, Th3844 presented more CAZymes with upregulated expression under cellulose degradation conditions than Th0179 and Ta0200, whereas a significant number of CAZymes from the last strain exhibited downregulated expression under such growth conditions (Fig. 5C). We would also like to highlight that important hemicellulases, such as GH10 and GH11, showed a notable difference in their expression profile between Ta0020 and Th3844. The same finding was observed for AA9 with the CBM1 domain (Fig. 5C). In contrast, CAZymes from the GT4 and GH20 families that exhibited upregulated expression under cellulose degradation conditions were identified for Ta0020 (Fig. 5C).

Moreover, the CAZyme families related to biomass degradation as well as mycoparasitism were investigated in greater depth (Fig. 6). In relation to lignocellulose depolymerization, CAZymes from the GH5, AA1, AA3, GH2, and GH3 families were well represented among all the strains, and the highest numbers were found in Th3844 and Th0179 (Fig. 6A). Regarding mycoparasitism activity, although CAZymes from the GH18 family, which are related to chitin degradation, were present in all evaluated strains, Tr0711 exhibited the smallest number of such enzymes in its genome. Additionally, although other CAZyme classes, which are related to mycoparasitic interactions, were present in the genomes of Th3844, Th0179, Ta0020, and Tr0711, the amount of each CAZyme family differed among the strains (Fig. 6B).

**Fig. 6.**
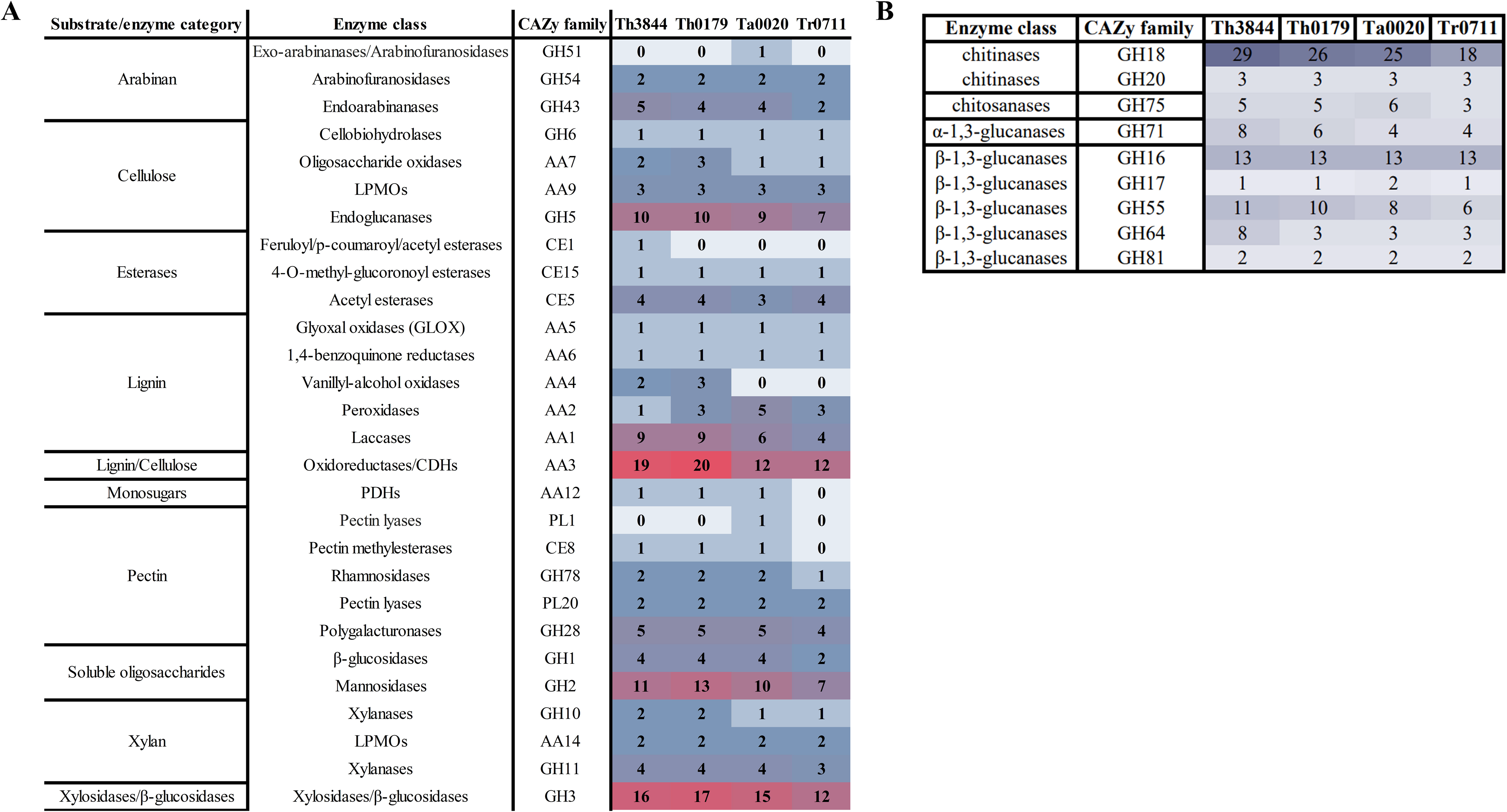
Quantitative comparison of the biomass-degrading and mycoparasitic enzymatic repertoires predicted for the *Trichoderma* isolate genomes. (a) Heatmap of the number of enzymes in each CAZy family from the Th3844, Th0179, Ta0020, and Tr0711 genomes. This map includes only the enzymes/proteins related to biomass degradation. (b) Heatmap of the number of enzymes in each CAZy family from the Th3844, Th0179, Ta0020, and Tr0711 genomes. This map includes only the enzymes related to mycoparasitic activity. Ta0020: *T. atroviride* CBMAI-0020; Th0179: *T. harzianum* CBMAI-0179; Th3844: *T. harzianum* IOC-3844; Tr0711: *T. reesei* CBMAI-0711; LPMOs: lytic polysaccharide monooxygenases; CDHs: cellobiose dehydrogenases; PHDs: pyranose dehydrogenases

### 3.4 Orthology analysis, phylogenetic profiling, and structural variant analyses

We identified a total of 17,188 orthogroups, which encompassed 236,336 genes (94.4%) in a total of 250,404 genes, i.e., the number of unassigned genes was equal to 14,068 (5.6%). Moreover, we detected 3,569 orthogroups among all present species, and 2,195 of these orthogroups consisted entirely of single-copy genes. Furthermore, 405 species-specific orthogroups, i.e., orthogroups that consist entirely of genes from one species, encompassing 1,066 genes (0.4%), were detected. Fifty percent of all genes were in orthogroups with 22 or more genes (G50 was 22) and were contained in the largest 4,252 orthogroups (O50 was 4,572) (Supplementary Material 5: Supplementary Table 11). The analysis of the orthologous relationships across the evaluated strains revealed that both *T. harzianum* strains shared the highest number of orthologous genes among them compared with the other strains. In relation to the other evaluated strains, Th3844 and Th0179 exhibited more orthologs in common with Ta0020 than with Tr0711 (Table 4). To explore the evolutionary history of Th3844, Th0179, Ta0020, and Tr0711, a rooted species tree was inferred using the 2,195 single-copy orthologous genes conserved in the 22 analyzed fungi (Fig. 7 and Supplementary Material 1: Supplementary Fig. 6). The phylogenetic analysis indicated that even though it was in the same clade as *T. reesei* QM6a, Tr0711 was most closely related to *Trichoderma parareesei* CBS 125925, while Ta0020 was most closely related to *T. gamsii* T6085. On the other hand, Th3844 was phylogenetically close to *T. harzianum* CBS 226.95 and *T. harzianum* TR274, while Th0179 was in the same clade as *T. guizhouense* NJAU 4742 and *T. afroharzianum* T6776.

**Fig. 7.**
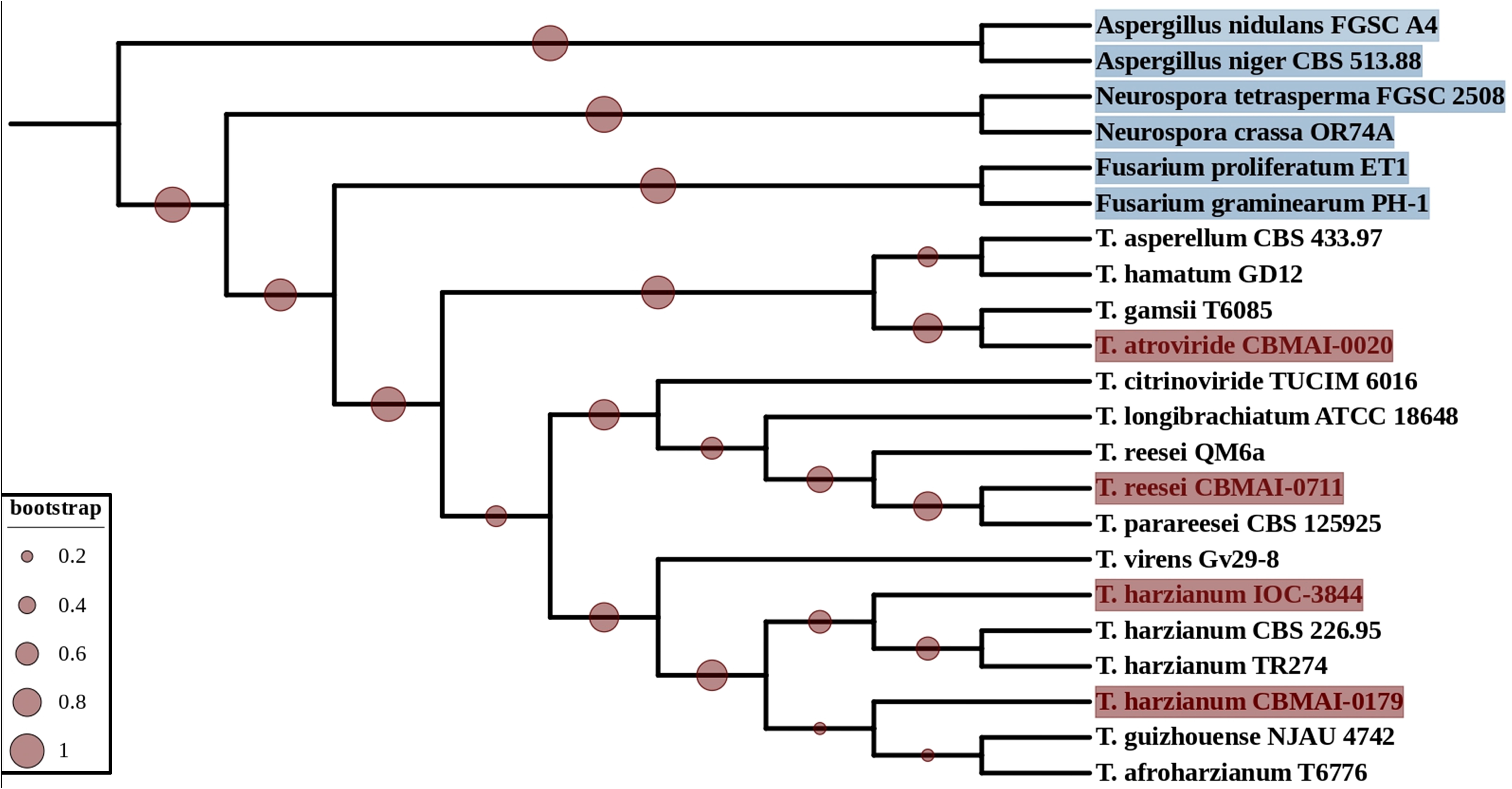
Phylogenetic relationships of *Trichoderma* spp. as inferred by an orthology analysis. The phylogenetic tree modeled by OrthoFinder software was based on the concatenation of 2,229 single-copy orthogroups. In addition to the proteomes of Th3844, Th0179, Ta0020, and Tr0711, this methodology shows the inferred relationships among 19 *Trichoderma* spp., for which the proteomes are available in the NCBI database. *Fusarium* spp., *Aspergillus* spp., and *Neurospora* spp. were used as the outgroup. Bootstrap values are shown at the nodes.

**Table 4.** Distribution of Th3844, Th0179, Ta0020, and Tr0711 orthologs.

|  | **Th3844** | **Th0179** | **Ta0020** | **Tr0711** |
| --- | --- | --- | --- | --- |
| **Th3844** | - | 9,933 | 8,273 | 7,572 |
| **Th0179** | 9,847 | - | 8,978 | 8,293 |
| **Ta0020** | 8,104 | 8,940 | - | 7,225 |
| **Tr0711** | 7,369 | 8,188 | 7,871 | - |

The SVs of the evaluated *Trichoderma* isolates were identified by mapping the long reads of the fungi against the reference genome of *T. reesei* QM6a (Martinez et al., 2008). A total of 12,407 (Th3844), 12,763 (Th0179), 11,650 (Ta0020), and 7,103 (Tr0711) SVs were identified for each strain, showing substitution rates of ∼1/2,674 nucleotides (Th3844), ∼1/2,585 nucleotides (Th0179), ∼1/2,832 nucleotides (Ta0020), and ∼1/4,655 nucleotides (Tr0711). These SVs included different phenomena that affect gene sequences, such as break ends, deletions, multiple nucleotides and InDels, duplications, insertions, and inversions (Supplementary Material 6: Supplementary Table 12 and Fig. 8A). Compared with the other evaluated strains, Tr0711 presented a low number of SVs, while Th0179 displayed the highest number. For all evaluated strains, the most frequently presented SV categories were multiple nucleotides and an InDel, followed by deletions and insertions (Fig. 8A).

**Fig. 8.**
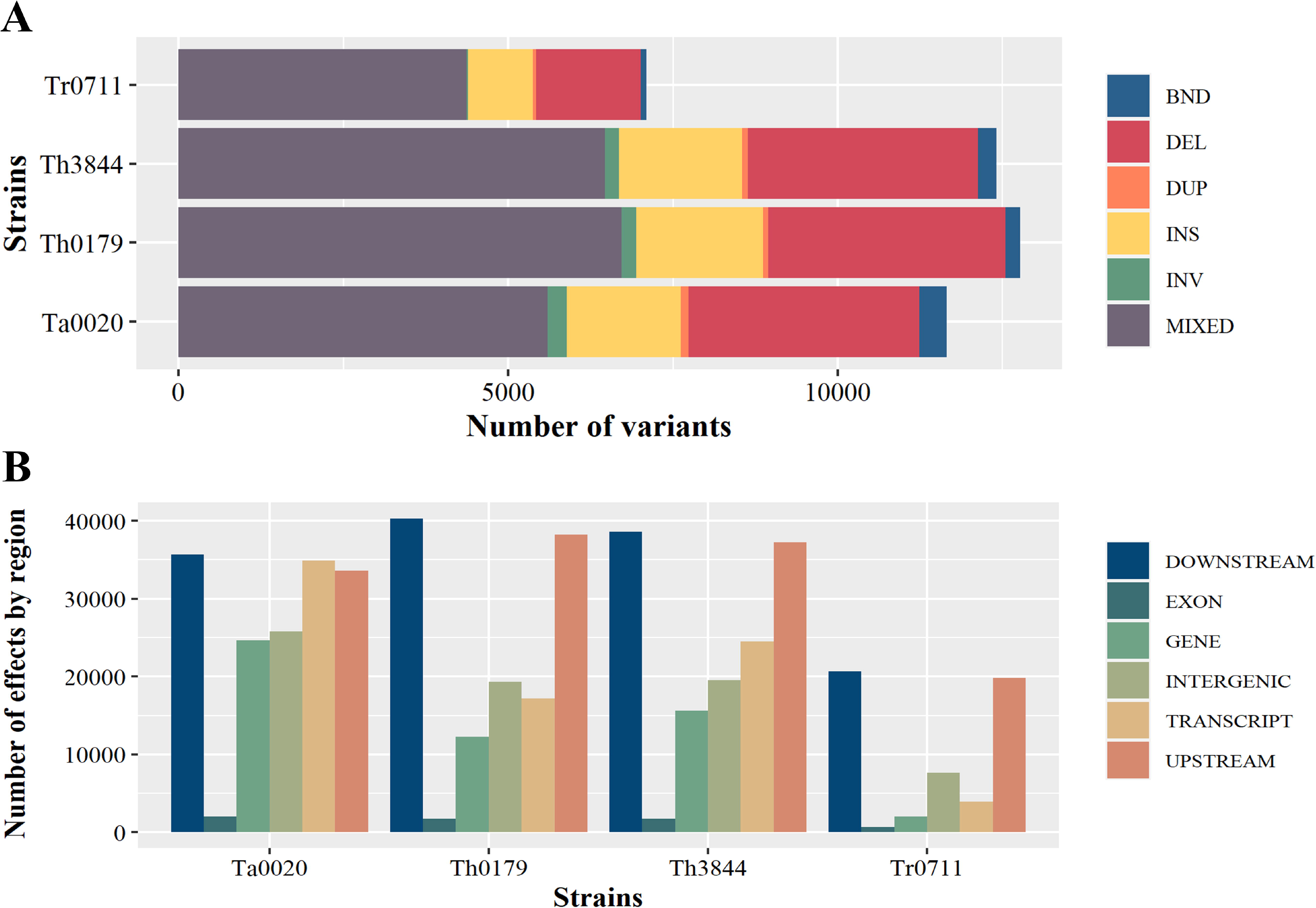
Structural variant heterogeneity across the genomes of the evaluated *Trichoderma* spp. (a) Long-read alignment-based structural variant (SV) analyses among the evaluated *Trichoderma* isolates and *T. reesei* QM6a showed breakends (BNDs), deletions (DELs), multiple nucleotides and InDels (MIXEDs), duplications (DUPs), insertions (INSs), and inversions (INVs) among the genomes. (B) Functional effects of the identified SVs. Ta0020: *T. atroviride* CBMAI-0020; Th0179: *T. harzianum* CBMAI-0179; Th3844: *T. harzianum* IOC-3844; Tr0711: *T. reesei* CBMAI-0711

To thoroughly investigate the functional effects of the identified SVs, we performed an annotation of the structural rearrangements, which were placed into different classes based on their predicted effects on protein function. Details of these effects can be found in Supplementary Material 7: Supplementary Tables 13-16, and the most prevalent effects are represented in Fig. 8B. For all evaluated strains, the majority of variants presented a modifier impact, which was higher at downstream and upstream genomic locations. Such an effect was more accentuated for both *T. harzianum* strains. In addition, SVs present in transcripts, genes, and intergenic regions were well represented for Ta0020.

With the aim of elucidating the SVs identified against the *T. reesei* QM6a reference genome in the evaluated genomes, we selected the CAZymes identified as DEGs under cellulose and glucose growth conditions and with orthologs in Th3844, Th0179, and Ta0020 for detailed analyses. After searching for corresponding homologs in the *T. reesei* QM6a reference genome, we investigated SVs in the following genes: GH72 (XM_006965422.1), GH1 (XM_006963374.1), GH20 (XM_006963001.1), GH16 (XM_006966546.1), GH1 (XM_006964776.1), and GH20 (XM_006969114.1). Considering such genes, we identified SVs in (I) 4 (Th3844), (II) 5 (Th0179), and (III) 4 (Ta0020) genes, and the majority had a modifier impact.

## 4 Discussion

Diversity within the genus *Trichoderma* is evident from the wide range of phenotypes exhibited by the fungi as well as the various ecological roles and industrial purposes the fungi serve (Nakkeeran et al., 2021). Because of their various applications, different *Trichoderma* species have become model organisms for a broad spectrum of physiological phenomena, such as plant cell wall degradation and enzyme production (Fang et al., 2021), biocontrol (Zin and Badaluddin, 2020), and response to light (Schmoll, 2018). Within the genus *Trichoderma*, *T. harzianum* has been used as a commercial biocontrol agent against plant diseases (Fraceto et al., 2018). In addition to their mycoparasitic activities, hydrolytic enzymes from *T. harzianum* strains have demonstrated great potential in the conversion of lignocellulosic biomass into fermentable sugars (Almeida et al., 2021; Delabona et al., 2020; Motta et al., 2021; Zhang et al., 2020). Recently, different types of enzymatic profiles across *Trichoderma* species were reported, in which Th3844 and Th0179 presented a higher hydrolytic potential during growth on cellulose than Ta0020 (Almeida et al., 2021); furthermore, differences between Th3844 and Th0179 concerning the transcriptional regulation coordinated by XYR1 and CRE1 during cellulose degradation were reported (Rosolen et al., 2022). Because such diversity in enzyme response might be related to transcriptomic and genomic differences, we aimed to provide foundations for further studies that could investigate such variations at a genomic level.

Herein, we presented high-quality genome assemblies for two *T. harzianum* strains with hydrolytic potential (Th3844 and Th0179) and, for comparative purposes, for a mycoparasitic species (Ta0020) and saprotrophic species (Tr0711). Thus, *T. reesei* and *T. atroviride* strains were used to assess the genetic differences in the genus *Trichoderma*. Except for Tr0711, the resulting genomic assemblies displayed the highest coverage scores and the lowest fragmentation values compared to those of *Trichoderma virens* Gv29-8 (Kubicek et al., 2011) as well as to *T. harzianum* T6776 (Baroncelli et al., 2015) and *T. atroviride* IMI206040 (Kubicek et al., 2011), which were used as the reference genomes in preceding studies from our research group (Almeida et al., 2021; Horta et al., 2018). Although the genomes were not assembled at the chromosome level, the quality of the Th3844, Th0179, Ta0020, and Tr0711 genome assemblies based on the BUSCO value was over 90%. Only the Th3844 genome exhibited 9.7% missing genes. However, even chromosome-level genome assembly does not necessarily achieve a complete BUSCO score (Chung et al., 2021).

After evaluating the quality of the four assembled genomes, we performed gene prediction and functional annotation for the datasets. The ecological behavior of the mycoparasites *T. atroviride* and *T. virens*, compared to the plant wall degrader *T. reesei*, is reflected by the sizes of the respective genomes; *T. atroviride* (36 Mb) and *T. virens* (39 Mb) were somewhat larger than the weakly mycoparasitic *T. reesei* (33 Mb) (Kubicek et al., 2011; Martinez et al., 2008). Herein, compared to Th3844 (40 Mb), Th0179 (39 Mb), and Ta0020 (36 Mb), the genome of Tr0711 (32 Mb) was smaller, which might be conceivably because the gene function was lost to mycoparasitism during the evolution of *T. reesei* (Kubicek et al., 2011). In relation to the number of genes, our results showed that Tr0711 presented a smaller gene content than Th3844, Th0179, and Ta0020. Such findings corroborated a previous study (Xie et al., 2014), in which 9,143 and 11,865 genes were predicted for *T. reesei* and *T. atroviride*, respectively. A recent general comparison of the genomes of twelve *Trichoderma* species revealed that their genetic repertoire varies in size (33-41□Mb) and in the number of predicted genes (9,292 and 14,095) (Kubicek et al., 2019a). In relation to the *T. reesei* QM6a reference genome, the genomes of Th3844, Th0179, Ta0020, and Tr0711 displayed significant structural reorganization, which was more greatly accentuated by an increased phylogenetic distance. Interestingly, this structural reorganization was also observed within strains of the same species, highlighting their genetic diversity.

### 4.1 Comparative and functional genomics

To obtain insights regarding the functional profile of Th3844, Th0179, Ta0020, and Tr0711, COG analyses of proteins from their genomes were performed. “Carbohydrate metabolism and transport” was a notable COG term for all evaluated strains, suggesting that the genomic arsenal of these fungi is connected to their ability to use carbon sources that are available in the environment. Such characteristics are well known for the saprophytic fungus *T. reesei* (Arntzen et al., 2020), and recent studies have observed the same characteristics for *T. harzianum* (Almeida et al., 2021; Delabona et al., 2020). “Posttranslational modification, protein turnover, chaperone functions” was the second most notable COG term that was present in the four evaluated genomes. Posttranslational modifications (PTMs), which are used by eukaryotic cells to diversify their protein functions and dynamically coordinate their signaling networks, encompass several specific chemical changes that occur on proteins following their synthesis (Ramazi and Zahiri, 2021).

The high extracellular secretion capability and eukaryotic PTM machinery make *Trichoderma* spp. particularly interesting hosts (Wei et al., 2021). In this context, PTMs are a major factor in the cellulolytic performance of fungal cellulases (Amore et al., 2017; Beckham et al., 2012; Dana et al., 2014), and the impact of plant PTMs on the enzyme performance and stability of the major cellobiohydrolase Cel7A from *T. reesei* has already been determined (van Eerde et al., 2020). In addition, PTMs, especially phosphorylation, of the proteins involved in plant biomass degradation, including CRE1, play an essential role in signal transduction to achieve carbon catabolite repression (CCR) (Han et al., 2020; Horta et al., 2019). Thus, describing this class of *Trichoderma* genomes is essential to understand the impact of alternative PTMs on the catalytic performance and stability of recombinant enzymes. We would also like to highlight that “secondary structure” and “amino acid transport and metabolism”, which are related to PTMs, were overrepresented COG terms.

SMs from microorganisms may play an antifungal role against agriculturally important phytopathogenic fungi (Khan et al., 2020a). Many *Trichoderma* species are the most prominent producers of SMs with antimicrobial activity against phytopathogenic fungi (Khan et al., 2020b; Morais et al., 2022). Therefore, we performed genome-wide identification and analysis of SMGCs in the Th3844, Th0179, Ta0020, and Tr0711 genomes. Considering the diversity of bioactive molecules isolated from the genus and given the vast biosynthetic potential that emerged from the antiSMASH analysis conducted in our study, the four evaluated strains, particularly Th0179, have high potential to produce bioactive molecules that warrant their use as biocontrol agents against plant pathogens. Furthermore, in both *T. harzianum* strains investigated in this study, namely, Th3844 and Th0179, genome mining identified the biosynthesis of tricholignan A, which is a natural product that helps plants assimilate iron from the soil (Chen et al., 2019). Similar results were reported for the biofertilizer strain *T. harzianum* t-22 (Chen et al., 2019). Detailed information about these SMs, when grouped together, enhances the understanding of their efficient utilization and allows further exploration of new bioactive compounds for the management of plant pathogenic fungi.

The comparative genomics of *Trichoderma* spp. suggested that mycoparasitic strains, such as *T. vires* and *T. atroviride*, presented a set of genes, including CAZymes and genes encoding SMs, that were more highly expressed and related to mycoparasitism than the saprotrophic species *T. reesei* (Kubicek et al., 2011). Although such fungi are widely used in industry as a source of cellulases and hemicellulases, they have a smaller arsenal of genes that encode the CAZymes related to biomass deconstruction than other lignocellulolytic fungi (Martinez et al., 2008). Regarding the CAZyme content, the results found here for the Th3844, Th0179, Ta0020, and Tr0711 genomes follow the same profile as that of a previous study (Fanelli et al., 2018), in which the CAZyme genetic endowment of some strains from *T. harzianum*, including B97 and T6776, was significantly higher than that of *T. atroviride* IMI206040, *T. reesei* QM6a, and *T. virens* Gv-29-8. However, by normalizing the CAZyme counts by the total gene counts for each strain, we found similar values among the evaluated fungi, as follows: (I) 3.8% (Th3844), (II) 3.6% (Th0179), (III) 3.7% (Ta0020), and 3.7% (Tr0711). Such results indicate that the probable differences regarding the CAZyme distribution might be related to the specific CAZyme families of a strain, as explored below, and not necessarily related to the total CAZyme content across the compared genomes. Furthermore, the presence of putative CAZyme-encoding genes in the genomes of Th0179 and Th3844 provides insight into their lignocellulose-degrading enzyme potential but cannot be directly related to their real degradation ability. In fact, since fungal species rely on different strategies, it has been observed that the number of genes related to the degradation of a given polysaccharide is not necessarily correlated to the extent of its degradation (Arntzen et al., 2020; Kjaerbolling et al., 2020). For this reason, CAZy analysis is associated with functional approaches, such as enzymatic activity assays, which provide valuable insight into the actual behavior of the concerned species on specific lignocellulose substrates.

In relation to the CAZyme families that are directly associated with the deconstruction of plant biomass, the genomes of Th3844, Th0179, Ta0020, and Tr0711 contained genes encoding GH5, which includes cellulases that are most highly abundant in fungi (Li and Walton, 2017); GH3, which includes β-glucosidases that are frequently secreted into the medium (Guo et al., 2016); and AA3, which is a member of the enzyme arsenal that is auxiliary to GHs (Levasseur et al., 2013). Lytic polysaccharide monooxygenases (LPMOs), which are classified into the CAZy auxiliary activity families AA9-AA11 and AA13-AA16, are copper-dependent enzymes that also perform important roles in lignocellulose degradation (Couturier et al., 2018; Monclaro and Filho, 2017). Herein, each of the Th3844, Th0179, Ta0020, and Tr0711 genomes exhibited three AA9 and two AA14 enzymes. Compared to other fungi, the genomes of *Trichoderma* species harbor a high number of chitinolytic genes (Kubicek et al., 2011, 2019), reflecting the importance of these enzymes in the mycoparasitic characteristics of fungi. From the *Trichoderma* genomes that have been analyzed in detail thus far, the fungal chitinases that belong to the family GH18 are significantly expanded in *T. virens*, *T. atroviride*, *T. harzianum*, *Trichoderma asperellum*, *Trichoderma gamsii*, and *Trichoderma atrobrunneum* (Fanelli et al., 2018; Kubicek et al., 2011). Similarly, the number of chitosanases (GH75) is enhanced, and there are at least five corresponding genes; in contrast, most other fungi have only one or two corresponding genes (Kappel et al., 2020; Kubicek et al., 2011). Furthermore, β-1,3-glucanases that belong to GH families 16, 17, 55, 64, and 81 are expanded in *Trichoderma* mycoparasites compared to other fungi (Fanelli et al., 2018; Kubicek et al., 2011). Here, the CAZyme families that are related to mycoparasitic activity were present in the four genomes studied, and most were in Th3844 and Th0179.

Transcriptome analysis identified the DEGs and the differentially expressed CAZyme genes in all strains. Although *T. reesei* is the primary source of hydrolytic enzymes that form the enzymatic cocktails, Tr0711 exhibited the smallest number of identified DEGs under cellulose growth conditions, whereas Th3844 presented the highest number, followed by Th0179 and Ta0020. Each strain presented a different set of genes with different expression levels, which can be attributed to between-strain differences in the regulatory mechanisms of hydrolysis.

### 4.2 Insights into evolutionary history

Kubicek et al. (Kubicek et al., 2019a) performed a time-scaled phylogenetic analysis using 638 orthologous genes of twelve species of fungi from the genus *Trichoderma*. Aiming to expand this analysis, we included the protein repertoire from the strains evaluated in this study, i.e., Th3844, Th0179, Tr0711, and Ta0020. The authors observed that the species *T. harzianum*, *T. reesei*, and *T. atroviride* are phylogenetically distant when compared to other fungi from the same genus, which we also observed. It is important to note that the bootstrap value observed by Kubicek et al. (Kubicek et al., 2019a) for *T. harzianum* strains was low when considering the maximum value as 100%. In this study, a low bootstrap value for the clade formed by aligning the sequence proteomes from Th0179, *T. guizhouense*, and *T. afroharzianum* was also observed. These observations might be explained by the complex speciation process within the *T. harzianum* species group (Druzhinina et al., 2010); therefore, the phylogenetic position is uncertain for strains of these fungal species. However, the molecular identification of Th3844 and Th0179 based on the ITS and *tef1* sequences has already been reported (Rosolen et al., 2022), which confirmed that both strains were phylogenetically close to other *T. harzianum* strains.

Overall, by applying orthology analyses, we identified orthogroups and orthologs between the evaluated strains, as well as among some other *Trichoderma* spp. and filamentous fungi that are more genetically distant. In the genus *Trichoderma*, several lifestyles have been documented, including saprotrophy, which is a lifestyle that is also observed in other filamentous fungi, such as *Neurospora* spp., *Aspergillus* spp., and *Fusarium* spp. (Arntzen et al., 2020; Correa et al., 2020; Najjarzadeh et al., 2021). Thus, the proteomes of some species and strains of such genera were also included in the orthology analysis. Through our results, we may infer that some genus-specific genes are necessary for specific lifestyles and are shared by fungi that have the same lifestyle but are in quite different evolutionary orders.

In this study, we detected SVs by aligning our four genomes against *T. reesei* QM6a (Martinez et al., 2008); although the genomes were assembled at a scaffolding level, unlike the *T. reesei* QM6a reference genome, which was assembled at the chromosome level, we opted to proceed with this dataset because its annotation file was available (Li et al., 2017). As expected, due to phylogenetic proximity to the reference genome (Rosolen et al., 2022), Tr0711 exhibited a lower number of SVs than the other strains. However, although *T. atroviride* is phylogenetically distant from *T. reesei* (Rosolen et al., 2022), Ta0020 exhibited fewer SVs than both *T. harzianum* strains, and this result might be explained by the uncertain phylogenetic position of fungi in these species (Druzhinina et al., 2010). Although the *T. harzianum* strains are phylogenetically close (Rosolen et al., 2022), comparison of the SVs identified from the mapping of both genomes against *T. reesei* QM6a reveals genetic variability across the strains (Martinez et al., 2008). Such results are consistent with the findings of previous studies in that genetic variations between fungal strains of the same species are not uncommon (Andersen et al., 2011; de Vries et al., 2017; Thanh et al., 2019).

Considering their basic and economic importance, the high-quality genomes found herein might be helpful for better understanding the diversity within the genus *Trichoderma*, as well as improving the biotechnological applications of such fungi. Furthermore, the comparative study of multiple related genomes might be helpful for understanding the evolution of genes that are related to economically important enzymes and for clarifying the evolutionary relationships related to protein function.

## Acknowledgments

We are grateful to CBMAI Campinas and SP for conceiving the fungal isolates used in the current study; the Center of Molecular Biology and Genetic Engineering (CBMEG) at the University of Campinas and SP for the use of the center and laboratory space; and the São Paulo Research Foundation (FAPESP), the Coordination of Improvement of Higher Education Personnel (CAPES, Computational Biology Program), and the Brazilian National Council for Technological and Scientific Development (CNPq) for supporting the project and researchers.

## Ethics approval

Not applicable.

## Informed consent

Not applicable.

## Data availability

All data generated or analyzed in this study are included in this published article (and its supplementary information files). The raw datasets and the assembled genomes were deposited in the NCBI Sequence Read Archive and can be accessed under BioProject number PRJNA781962 and BioSample number as follows: SAMN23309297 (Th3844), SAMN23309298 (Th0179), SAMN23309299 (Ta0020), and SAMN23309300 (Tr0711).

## Authors’ contribution

**RRR:** Writing – original draft, methodology, formal analysis, and visualization. **MACH:** Conceptualization, methodology, and writing – review & editing. **PHCA:** Methodology and resources. **CCS:** Methodology and formal analysis. **DAS:** Methodology and resources. **GHG:** Writing - review & editing. **APS:** Conceptualization, supervision, review & editing, and funding acquisition.

