## Supplementary Material 1 for "Whole-Genome Sequencing and Comparative Genomic Analysis of Potential Biotechnological Strains of Trichoderma harzianum, Trichoderma atroviride*, and* Trichoderma reesei"

### Materials and methods

#### *Phylogenetic analyses*

The *rbp2* and *tef1* nucleotide sequences of *Trichoderma* spp. were retrieved from the NCBI database (https://www.ncbi.nlm.nih.gov/). Additionally, *tef1* nucleotide sequences amplified from the genomic DNA of Th3844, Th0179, Ta0020, and Tr0711 by PCR were kindly provided by the CBMAI and included in the phylogenetic analysis. These strains were used as reference genomes to retrieve the *rbp2* nucleotide sequences belonging to *Trichoderma* spp. from the NCBI database. Additionally, the *T. reesei* QM6a genome was used as a reference genome to retrieve the *rbp2* nucleotide sequences belonging to *Trichoderma* spp. from the NCBI database and to identify the corresponding homologs of Th3844, Th0179, Ta0020, and Tr0711 by using BLASTn. These sequences were included in the phylogenetic analyses. For *rpb2*, *Fusarium* spp. and *Neurospora* spp. were used as outgroups.

Multiple sequence alignment was performed using ClustalW (Thompson et al., 1994), and a phylogenetic tree was created using Molecular Evolutionary Genetics Analysis (MEGA) software v7.0 (Kumar et al., 2016). For *rbp2* and *tef1*, the maximum likelihood (ML) (Jones et al., 1992) method of inference was used based on a (I) Tamura 3-parameter (T92) model (Tamura and Nei, 1993) and a (II) Hasegawa-Kishino-Yano (HKY) model (Hasegawa et al., 1985), respectively. We included 1,000 bootstrap replicates (Felsenstein, 1985) in each analysis, and the trees were visualized and edited using Interactive Tree of Life (iTOL) v6 (https://itol.embl.de/).

### Results

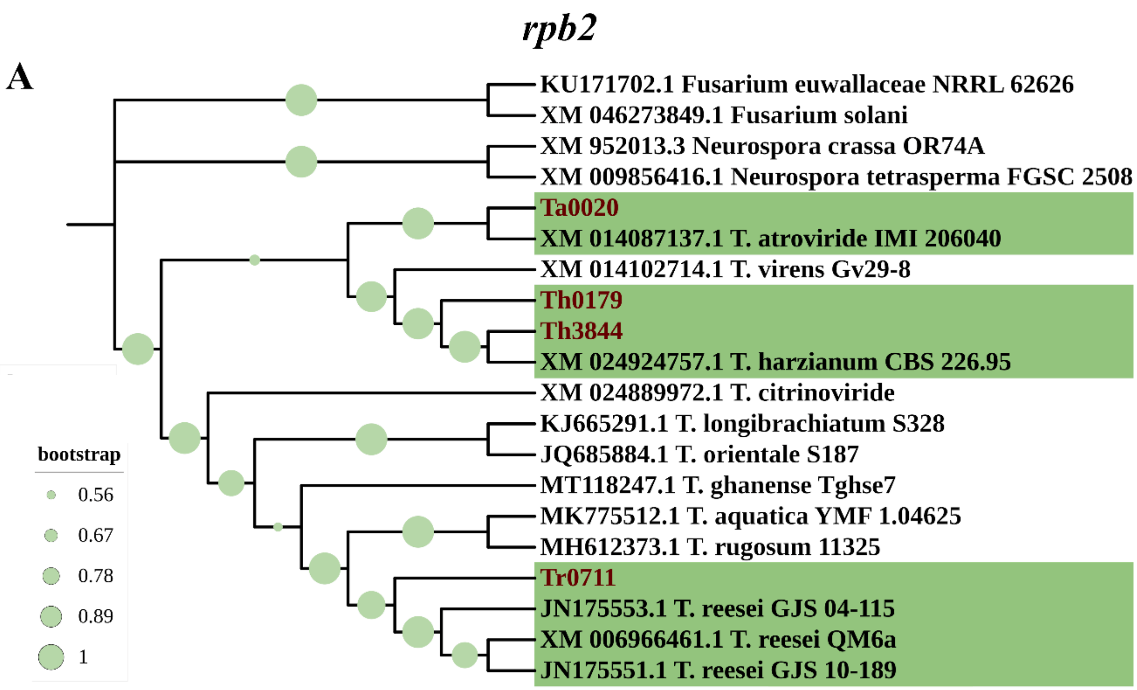

rpb2
A
KU171702.1 Fusarium euwallaceae NRRL 62626
XM 046273849.1 Fusarium solani
XM 952013.3 Neurospora crassa OR74A
XM 009856416.1 Neurospora tetrasperma FGSC 2508
Ta0020
XM 014087137.1 T. atroviride IMI 206040
XM 014102714.1 T. virens Gv29-8
Th0179
Th3844
XM 024924757.1 T. harzianum CBS 226.95
XM 024889972.1 T. citrinoviride
KJ665291.1 T. longibrachiatum S328
JQ685884.1 T. orientale S187
MT118247.1 T. ghanense Tghse7
MK775512.1 T. aquatica YMF 1.04625
MH612373.1 T. rugosum 11325
Tr0711
JN175553.1 T. reesei GJS 04-115
XM 006966461.1 T. reesei QM6a
JN175551.1 T. reesei GJS 10-189
bootstrap
0.56
0.67
0.78
0.89
1


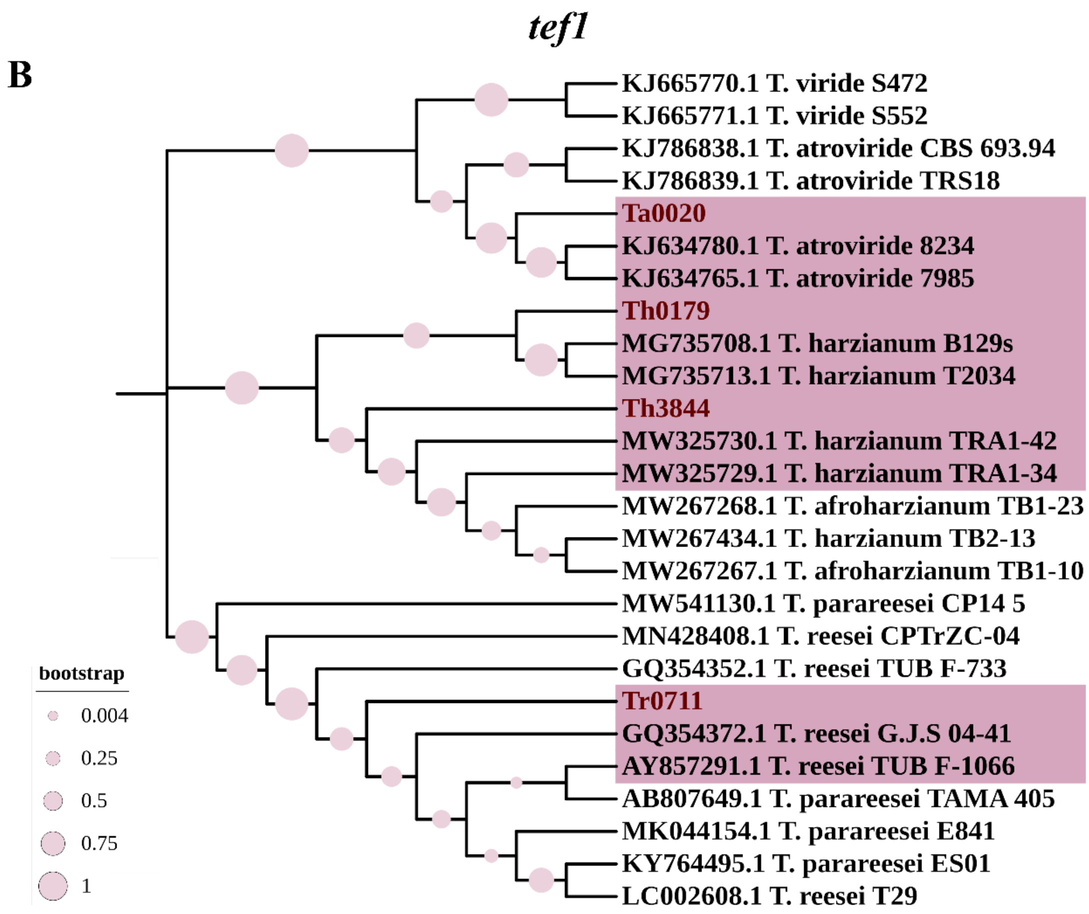

tef1
B
KJ665770.1 T. viride S472
KJ665771.1 T. viride S552
KJ786838.1 T. atroviride CBS 693.94
KJ786839.1 T. atroviride TRS18
Ta0020
KJ634780.1 T. atroviride 8234
KJ634765.1 T. atroviride 7985
Th0179
MG735708.1 T. harzianum B129s
MG735713.1 T. harzianum T2034
Th3844
MW325730.1 T. harzianum TRA1-42
MW325729.1 T. harzianum TRA1-34
MW267268.1 T. afroharzianum TB1-23
MW267434.1 T. harzianum TB2-13
MW267267.1 T. afroharzianum TB1-10
MW541130.1 T. parareesei CP14 5
MN428408.1 T. reesei CPTrZC-04
GQ354352.1 T. reesei TUB F-733
Tr0711
GQ354372.1 T. reesei G.J.S 04-41
AY857291.1 T. reesei TUB F-1066
AB807649.1 T. parareesei TAMA 405
MK044154.1 T. parareesei E841
KY764495.1 T. parareesei ES01
LC002608.1 T. reesei T29
bootstrap
0.004
0.25
0.5
0.75
1

**Supplementary Material 1: Supplementary Figure 1.** Phylogenetic relationships of *Trichoderma* spp. inferred through the analysis of *rpb2* and *tef1* nucleotide sequences. Sequences corresponding to the *rpb2* (A) and *tef1* (B) nucleotide sequences were used to analyze and compare the phylogenetic relationships of the studied *Trichoderma* species/strains. By using BLASTn, the *rpb2* sequences of Th3844, Th0179, Ta0020, and Tr0711 were recovered from the assembled genomes using the *T. reesei* QM6a *rpb2* sequence as the reference genome. The *tef1* sequences of Th3844, Th0179, Ta0020, and Tr0711 were amplified from genomic DNA. The other *tef1* and *rpb2* sequences were derived from the NCBI database. Th3844: *T. harzianum* IOC-3844; Th0179: *T. harzianum* CBMAI-0179; Ta0020: *T. atroviride* CBMAI-0020; Tr0711: *T. reesei* CBMAI-0711.

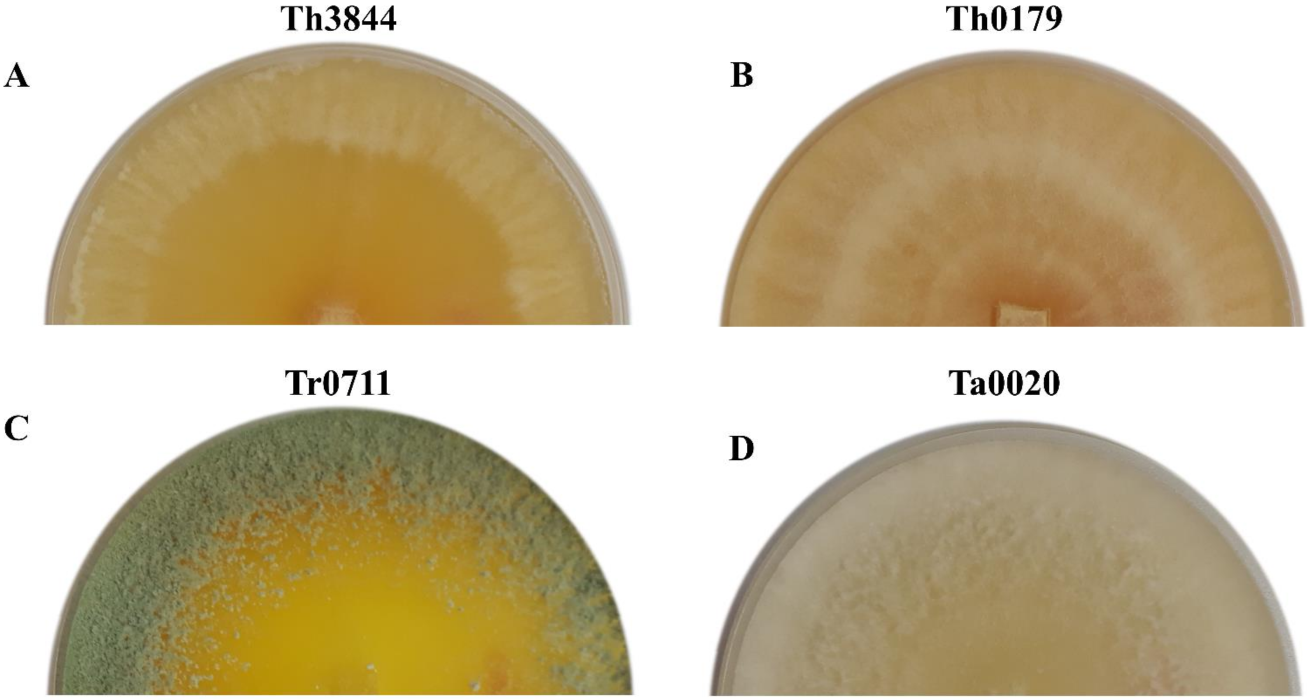


**Supplementary Material 1: Supplementary Figure 2. Physical morphology of the strains evaluated in this study.** (A) Th3844, (B) Th0179, (C) Ta0020, and (D) Tr0711 were cultivated on potato dextrose agar (PDA) solid medium (100 µg/ml ampicillin and 34 µg/ml chloramphenicol) for 3 days at 28 °C. Although the four strains showed similar radial growth, these fungi presented different color patterns; i.e., both strains of *T. harzianum* predominantly exhibited a yellow color, Tr0711 showed a green color, and Ta0020 presented a white color. Th3844: *T. harzianum* IOC-3844; Th0179: *T. harzianum* CBMAI-0179; Ta0020: *T. atroviride* CBMAI-0020; Tr0711: *T. reesei* CBMAI-0711.

**Supplementary Material 1: Supplementary Table 1. Evaluation of gDNA quality.** The quality of the gDNA extracted from the Th3844, Th0179, Ta0020, and Tr0711 strains was evaluated using a NanoDrop 1000 spectrophotometer (Thermo Fisher Scientific) and Qubit Fluorometer (Thermo Fisher Scientific). The concentrations are given in ng/µl.

| Sample | Qubit-ug | 260/280 | 260/230 | Amount of water used for rehydration µL | Final concentration ng/µL |
|---|---|---|---|---|---|
| **Sample 1 - Ta0020 Speed vacuum** | 20 | 1.75 | 1.87 | 100 | 200 |
| **Sample 2 - Th0179 Speed vacuum** | 20 | 1.87 | 1.92 | 100 | 200 |
| **Sample 3 - Tr0711 Speed vacuum** | 22 | 1.93 | 1.98 | 100 | 220 |
| **Sample 4 - Th3844 Speed vacuum** | 22 | 1.85 | 1.82 | 100 | 220 |
| **Sample 5 - Ta0020 Precipitation stopped in the ETOH 70% step** | 21 | 1.84 | 1.89 | 100 | 210 |
| **Sample 6 - Th0179 Precipitation stopped in the ETOH 70% step** | 20 | 1.99 | 1.84 | 100 | 200 |
| **Sample 7 - Tr0711 Precipitation stopped in the ETOH 70% step** | 27 | 1.94 | 1.96 | 100 | 270 |
| **Sample 8 - Th3844 Precipitation stopped in the ETOH 70% step** | 23 | 2.01 | 1.85 | 100 | 230 |

Ta0020: *T. atroviride* CBMAI-0020; Th0179: *T. harzianum* CBMAI-0179; Th3844: *T. harzianum* IOC-3844; Tr0711: *T. reesei* CBMAI-0711.

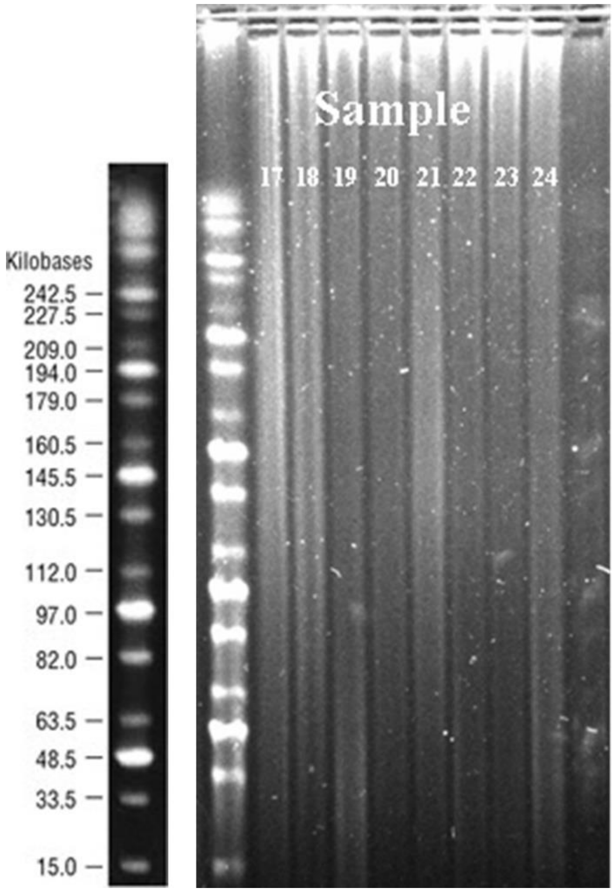


**Supplementary Material 1: Supplementary Figure 3. Evaluation of gDNA integrity on a 6% agarose gel.** Sample 17: Ta0020 (speed vacuumed); Sample 18: Th0179 (speed vacuumed); Sample 19: Tr0711 (speed vacuumed); Sample 20: Th3844 (speed vacuumed); Sample 21: Ta0020 (precipitated); Sample 22: Th0179 (precipitated); Sample 23: Tr0711 (precipitated); Sample 24: Th3844 (precipitated); Ta0020: *T. atroviride* CBMAI-0020; Th0179: *T. harzianum* CBMAI-0179; Th3844: *T. harzianum* IOC-3844; Tr0711: *T. reesei* CBMAI-0711.

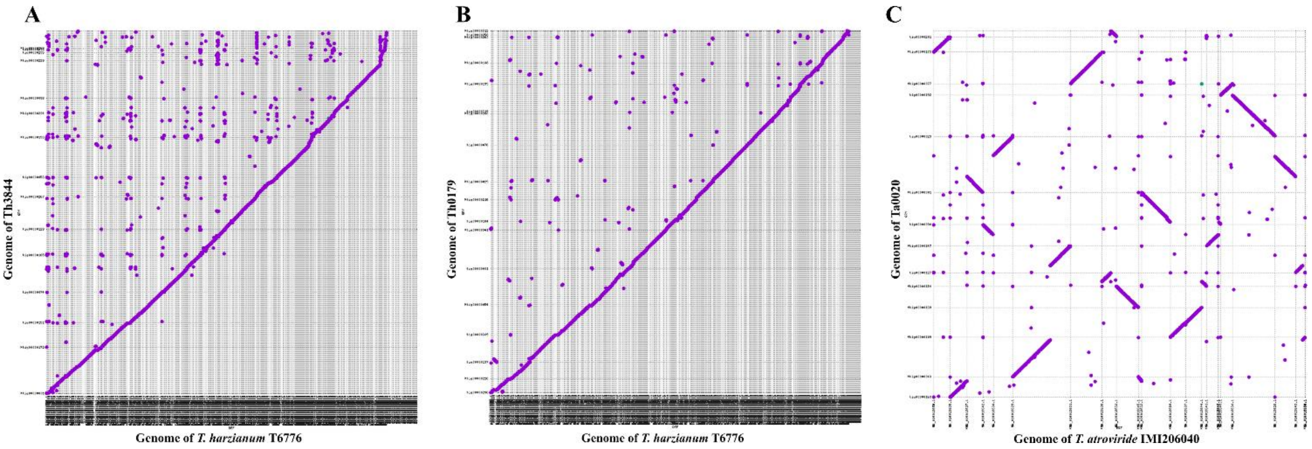


**Supplementary Material 1: Supplementary Figure 4. Comparisons between the genomes of the analyzed *Trichoderma* isolates and their respective reference genomes.** Dot plots of the assemblies of (**A**) Th3844, (**B**) Th0179, and (**C**) Ta0020 generated by Canu (y-axis) against those of *T. harzianum* T6776 (x-axis) and *T. atroviride* IMI206040 (x-axis) available in the NCBI database. Ta0020: *T. atroviride* CBMAI-0020; Th0179: *T. harzianum* CBMAI-0179; Th3844: *T. harzianum* IOC-3844.

**Supplementary Material 1: Supplementary Table 2.** Distribution of COG functional categories in *Trichoderma* spp. Each number in parentheses is the relative abundance.

| | | Th3844 | | Th0179 | | Ta0020 | | Tr0711 | |
|---|---|---|---|---|---|---|---|---|---|
| **COG class** | **COG functional categories** | **Count** | **(%)** | **Count** | **(%)** | **Count** | **(%)** | **Count** | **(%)** |
| **A** | **RNA processing and modification** | 279 | 3,7 | 307 | 3,7 | 305 | 3,8 | 300 | 4,1 |
| **B** | **Chromatin structure and dynamics** | 174 | 2,3 | 191 | 2,3 | 172 | 2,1 | 164 | 2,2 |
| **C** | **Energy production and conversion** | 318 | 4,2 | 338 | 4,1 | 324 | 4,0 | 285 | 3,9 |
| **D** | **Cell cycle control and mitosis** | 143 | 1,9 | 161 | 1,9 | 160 | 2,0 | 155 | 2,1 |
| **E** | **Amino acid metabolism and transport** | 420 | 5,5 | 449 | 5,4 | 411 | 5,1 | 389 | 5,3 |
| **F** | **Nucleotide metabolism and transport** | 104 | 1,4 | 118 | 1,4 | 110 | 1,4 | 109 | 1,5 |
| **G** | **Carbohydrate metabolism and transport** | 516 | 6,8 | 561 | 6,8 | 536 | 6,7 | 456 | 6,2 |
| **H** | **Coenzyme metabolism** | 177 | 2,3 | 197 | 2,4 | 191 | 2,4 | 175 | 2,4 |
| **I** | **Lipid metabolism** | 309 | 4,1 | 333 | 4,0 | 324 | 4,0 | 293 | 4,0 |
| **J** | **Translation** | 321 | 4,2 | 358 | 4,3 | 354 | 4,4 | 335 | 4,5 |
| **K** | **Transcription** | 284 | 3,7 | 312 | 3,8 | 305 | 3,8 | 301 | 4,1 |
| **L** | **Replication and repair** | 209 | 2,8 | 229 | 2,8 | 233 | 2,9 | 220 | 3,0 |
| **M** | **Cell wall/membrane/envelop biogenesis** | 92 | 1,2 | 101 | 1,2 | 110 | 1,4 | 81 | 1,1 |
| **N** | **Cell motility** | 4 | 0,1 | 5 | 0,1 | 5 | 0,1 | 5 | 0,1 |
| **O** | **Post-translational modification, protein turnover, chaperones** | 502 | 6,6 | 541 | 6,5 | 529 | 6,6 | 510 | 6,9 |
| **P** | **Inorganic ion transport and metabolism** | 229 | 3,0 | 251 | 3,0 | 242 | 3,0 | 221 | 3,0 |
| **Q** | **Secondary structure** | 422 | 5,6 | 473 | 5,7 | 425 | 5,3 | 353 | 4,8 |
| **S** | **Function unknown** | 2184 | 28,8 | 2402 | 28,9 | 2298 | 28,7 | 2107 | 28,5 |
| **T** | **Signal transduction** | 326 | 4,3 | 337 | 4,1 | 343 | 4,3 | 324 | 4,4 |
| **U** | **Intracellular trafficking, secretion, and vesicular transport** | 392 | 5,2 | 441 | 5,3 | 441 | 5,5 | 418 | 5,7 |
| **V** | **Defense mechanisms** | 51 | 0,7 | 54 | 0,6 | 48 | 0,6 | 35 | 0,5 |
| **W** | **Extracellular structures** | 5 | 0,1 | 5 | 0,1 | 4 | 0,0 | 5 | 0,1 |
| **Y** | **Nuclear structure** | 21 | 0,3 | 26 | 0,3 | 27 | 0,3 | 26 | 0,4 |
| **Z** | **Cytoskeleton** | 108 | 1,4 | 121 | 1,5 | 119 | 1,5 | 115 | 1,6 |

| | | | | | | | | | |
|---|---|---|---|---|---|---|---|---|---|
| **Total** | | 7590 | 100,0 | 8311 | 100,0 | 8016 | 100,0 | 7382 | 100 |

COG: Clusters of Orthologous Groups of proteins; Ta0020: *T. atroviride* CBMAI-0020; Th0179: *T. harzianum* CBMAI-0179; Th3844: *T. harzianum* IOC-3844; Tr0711: *T. reesei* CBMAI-0711.

**Supplementary Material 1: Supplementary Table 6.** Expression profiles of CAZymes in the genome of Th3844 that were differentially expressed under cellulose growth conditions. Transcript expression levels were calculated as TPM values. Genes with p values ≤ 0.05 and fold changes ≥ 1.5 (upregulated) or ≤ -1.5 (downregulated) were considered differentially expressed. Th3844: *T. harzianum* IOC-3844; T6776: *T. harzianum* T6776; FC values: fold change values; DEGs: differentially expressed genes.

| Gene ID_Th3844 | Protein product_ThT6776 | GeneID_ThT6776 | FC values | p value | DEG | Family | Secreted |
|---|---|---|---|---|---|---|---|
| **Th3844_004197** | lcl\|JOKZ01000169.1_cds_KKP02016.1_5897 | THAR02_05896 | 9,11 | 0,000 | Up | GH11 | Y |
| **Th3844_007490** | lcl\|JOKZ01000505.1_cds_KKO97781.1_10113 | THAR02_10113 | 7,18 | 0,000 | Up | AA9 | Y |
| **Th3844_001507** | lcl\|JOKZ01000041.1_cds_KKP05759.1_2134 | THAR02_02133 | 6,57 | 0,000 | Up | CBM1 | Y |
| **Th3844_001480** | lcl\|JOKZ01000041.1_cds_KKP05778.1_2153 | THAR02_02152 | 5,36 | 0,002 | Up | CBM42+GH54 | Y |
| **Th3844_001288** | lcl\|JOKZ01000027.1_cds_KKP06353.1_1502 | THAR02_01501 | 5,30 | 0,000 | Up | GH12 | Y |
| **Th3844_008264** | lcl\|JOKZ01000071.1_cds_KKP04658.1_3272 | THAR02_03271 | 5,30 | 0,000 | Up | GH106 | N |
| **Th3844_001506** | lcl\|JOKZ01000041.1_cds_KKP05760.1_2135 | THAR02_02134 | 5,23 | 0,000 | Up | AA9+CBM1 | N |

| | | | | | | | |
|---|---|---|---|---|---|---|---|
| **Th3844_004134** | lcl\|JOKZ01000798.1_cds_KKO96621.1_11277 | THAR02_11277 | 5,22 | 0,000 | Up | GH30_7 | Y |
| **Th3844_009080** | lcl\|JOKZ01000342.1_cds_KKO99257.1_8631 | THAR02_08630 | 5,22 | 0,003 | Up | GH11 | Y |
| **Th3844_009285** | lcl\|JOKZ01000266.1_cds_KKP00250.1_7664 | THAR02_07663 | 4,99 | 0,001 | Up | CE5 | Y |
| **Th3844_002706** | lcl\|JOKZ01000020.1_cds_KKP06706.1_1196 | THAR02_01195 | 4,58 | 0,048 | Up | GH128 | Y |
| **Th3844_008287** | lcl\|JOKZ01000328.1_cds_KKO99424.1_8480 | THAR02_08479 | 4,54 | 0,000 | Up | CBM1 | N |
| **Th3844_009686** | lcl\|JOKZ01000303.1_cds_KKO99737.1_8168 | THAR02_08167 | 4,05 | 0,000 | Up | CE5 | Y |
| **Th3844_002808** | lcl\|JOKZ01000353.1_cds_KKO99128.1_8763 | THAR02_08762 | 4,03 | 0,000 | Up | CBM1+CE0 | N |
| **Th3844_010327** | lcl\|JOKZ01000108.1_cds_KKP03494.1_4415 | THAR02_04414 | 3,92 | 0,000 | Up | CBM1+GH6 | Y |
| **Th3844_008201** | lcl\|JOKZ01000108.1_cds_KKP03485.1_4406 | THAR02_04405 | 3,85 | 0,000 | Up | CBM1+GH5_5 | Y |
| **Th3844_008923** | lcl\|JOKZ01000074.1_cds_KKP04531.1_3358 | THAR02_03357 | 3,81 | 0,000 | Up | CBM1+GH7 | Y |
| **Th3844_004135** | lcl\|JOKZ01000798.1_cds_KKO96622.1_11278 | THAR02_11278 | 3,73 | 0,000 | Up | CBM1+CE15 | Y |
| **Th3844_000586** | lcl\|JOKZ01000089.1_cds_KKP04060.1_3853 | THAR02_03852 | 3,60 | 0,005 | Up | CE8 | Y |
| **Th3844_010887** | lcl\|JOKZ01000185.1_cds_KKP01661.1_6251 | THAR02_06250 | 3,60 | 0,000 | Up | GH62 | Y |
| **Th3844_009378** | lcl\|JOKZ01000362.1_cds_KKO99033.1_8859 | THAR02_08858 | 3,52 | 0,000 | Up | GH11 | Y |
| **Th3844_001431** | lcl\|JOKZ01000724.1_cds_KKO96812.1_11082 | THAR02_11082 | 3,47 | 0,022 | Up | AA1_2 | Y |
| **Th3844_008286** | lcl\|JOKZ01000328.1_cds_KKO99423.1_8479 | THAR02_08478 | 3,46 | 0,001 | Up | GH30_7 | N |

| | | | | | | | |
|---|---|---|---|---|---|---|---|
| **Th3844_008290** | lcl\|JOKZ01000365.1_cds_KKO99004.1_8898 | THAR02_08897 | 3,42 | 0,000 | Up | CBM1+GH7 | Y |
| **Th3844_001491** | lcl\|JOKZ01000041.1_cds_KKP05773.1_2148 | THAR02_02147 | 3,30 | 0,000 | Up | GH11 | Y |
| **Th3844_006069** | lcl\|JOKZ01000450.1_cds_KKO98175.1_9720 | THAR02_09719 | 3,22 | 0,000 | Up | CBM1+GH5_5 | Y |
| **Th3844_000587** | lcl\|JOKZ01000089.1_cds_KKP04059.1_3852 | THAR02_03851 | 3,19 | 0,000 | Up | CBM1+GH5_7 | Y |
| **Th3844_004107** | lcl\|JOKZ01000064.1_cds_KKP04907.1_3009 | THAR02_03008 | 3,11 | 0,006 | Up | GH55 | Y |
| **Th3844_002606** | lcl\|JOKZ01000014.1_cds_KKP07011.1_891 | THAR02_00890 | 2,99 | 0,000 | Up | GH3 | Y |
| **Th3844_009872** | lcl\|JOKZ01000247.1_cds_KKP00524.1_7365 | THAR02_07364 | 2,98 | 0,002 | Up | GH55 | N |
| **Th3844_002607** | lcl\|JOKZ01000014.1_cds_KKP07012.1_892 | THAR02_00891 | 2,56 | 0,000 | Up | GH3 | Y |
| **Th3844_006588** | lcl\|JOKZ01000106.1_cds_KKP03537.1_4345 | THAR02_04344 | 2,52 | 0,023 | Up | CBM24+GH71 | Y |
| **Th3844_008024** | lcl\|JOKZ01000008.1_cds_KKP07377.1_575 | THAR02_00574 | 2,48 | 0,042 | Up | GH93 | Y |
| **Th3844_001697** | lcl\|JOKZ01000010.1_cds_KKP07226.1_657 | THAR02_00656 | 2,39 | 0,006 | Up | GH3 | Y |
| **Th3844_004936** | lcl\|JOKZ01000032.1_cds_KKP06161.1_1765 | THAR02_01764 | 2,28 | 0,010 | Up | GH30_5 | Y |
| **Th3844_008511** | lcl\|JOKZ01000017.1_cds_KKP06872.1_1070 | THAR02_01069 | 2,13 | 0,000 | Up | GH64 | Y |
| **Th3844_010527** | lcl\|JOKZ01000017.1_cds_KKP06872.1_1070 | THAR02_01069 | 2,13 | 0,000 | Up | GH64 | Y |
| **Th3844_010552** | lcl\|JOKZ01000017.1_cds_KKP06872.1_1070 | THAR02_01069 | 2,13 | 0,000 | Up | GH64 | Y |
| **Th3844_011449** | lcl\|JOKZ01000017.1_cds_KKP06872.1_1070 | THAR02_01069 | 2,13 | 0,000 | Up | GH64 | Y |
| **Th3844_011205** | lcl\|JOKZ01000259.1_cds_KKP00338.1_7550 | THAR02_07549 | 2,10 | 0,003 | Up | GH30_3 | Y |

| | | | | | | | |
|---|---|---|---|---|---|---|---|
| **Th3844_001038** | lcl\|JOKZ01000037.1_cds_KKP05943.1_1983 | THAR02_01982 | 1,99 | 0,044 | Up | GH35 | N |
| **Th3844_010884** | lcl\|JOKZ01000185.1_cds_KKP01663.1_6253 | THAR02_06252 | 1,97 | 0,000 | Up | CBM24+GH71 | Y |
| **Th3844_003274** | lcl\|JOKZ01000095.1_cds_KKP03872.1_4022 | THAR02_04021 | 1,95 | 0,000 | Up | GH72 | Y |
| **Th3844_007252** | lcl\|JOKZ01000044.1_cds_KKP05610.1_2252 | THAR02_02251 | 1,93 | 0,000 | Up | GH1 | N |
| **Th3844_008414** | lcl\|JOKZ01000128.1_cds_KKP02981.1_4913 | THAR02_04912 | 1,93 | 0,000 | Up | CBM18+GH18 | N |
| **Th3844_002223** | lcl\|JOKZ01000420.1_cds_KKO98433.1_9461 | THAR02_09460 | 1,84 | 0,000 | Up | GH71 | N |
| **Th3844_006635** | lcl\|JOKZ01000072.1_cds_KKP04612.1_3303 | THAR02_03302 | 1,83 | 0,000 | Up | GH16_3 | Y |
| **Th3844_003759** | lcl\|JOKZ01000149.1_cds_KKP02477.1_5433 | THAR02_05432 | 1,79 | 0,000 | Up | GH1 | N |
| **Th3844_009453** | lcl\|JOKZ01000309.1_cds_KKO99651.1_8236 | THAR02_08235 | 1,77 | 0,000 | Up | GH18 | Y |
| **Th3844_005916** | lcl\|JOKZ01000566.1_cds_KKO97447.1_10453 | THAR02_10453 | 1,73 | 0,000 | Up | CE5 | Y |
| **Th3844_004436** | lcl\|JOKZ01000087.1_cds_KKP04131.1_3791 | THAR02_03790 | 1,69 | 0,003 | Up | GH76 | Y |
| **Th3844_000059** | lcl\|JOKZ01000182.1_cds_KKP01722.1_6182 | THAR02_06181 | 1,67 | 0,014 | Up | GH18 | N |
| **Th3844_001563** | lcl\|JOKZ01000023.1_cds_KKP06557.1_1349 | THAR02_01348 | 1,63 | 0,001 | Up | GH106 | N |
| **Th3844_005244** | lcl\|JOKZ01000104.1_cds_KKP03616.1_4301 | THAR02_04300 | 1,55 | 0,034 | Up | GH2 | N |
| **Th3844_005513** | lcl\|JOKZ01000067.1_cds_KKP04790.1_3128 | THAR02_03127 | 1,52 | 0,038 | Up | AA11 | Y |

**Supplementary Material 1: Supplementary Table 7.** Expression profiles of CAZymes in the genome of Th0179 that were differentially expressed under cellulose growth conditions. Transcript expression levels were calculated as TPM values. Genes with p values ≤ 0.05 and fold changes ≥ 1.5 (upregulated) or ≤ -1.5 (downregulated) were considered differentially expressed. Th0179: *T. harzianum* CBMAI-0179; T6776: *T. harzianum* T6776; FC values: fold change values; DEGs: differentially expressed genes.

| Gene ID_Th0179 | Protein product_ThT6776 | GeneID_ThT6776 | FC values | p value | DEG | Family | Secreted |
|---|---|---|---|---|---|---|---|
| **Th0179_010397** | lcl\|JOKZ01000041.1_cds_KKP05758.1_2133 | THAR02_02132 | 2,893 | 0,037 | Up | GH3 | Y |
| **Th0179_003380** | lcl\|JOKZ01000149.1_cds_KKP02477.1_5433 | THAR02_05432 | 2,637 | 0,000 | Up | GH1 | N |
| **Th0179_010399** | lcl\|JOKZ01000041.1_cds_KKP05760.1_2135 | THAR02_02134 | 2,600 | 0,000 | Up | AA9+CBM1 | N |
| **Th0179_010293** | lcl\|JOKZ01000289.1_cds_KKO99924.1_7959 | THAR02_07958 | 2,598 | 0,023 | Up | GH27 | N |
| **Th0179_010398** | lcl\|JOKZ01000041.1_cds_KKP05759.1_2134 | THAR02_02133 | 2,273 | 0,003 | Up | CBM1 | Y |
| **Th0179_009135** | lcl\|JOKZ01000071.1_cds_KKP04658.1_3272 | THAR02_03271 | 2,251 | 0,003 | Up | GH10 | N |
| **Th0179_004888** | lcl\|JOKZ01000044.1_cds_KKP05610.1_2252 | THAR02_02251 | 2,215 | 0,000 | Up | GH1 | N |
| **Th0179_004051** | lcl\|JOKZ01000056.1_cds_KKP05192.1_2712 | THAR02_02711 | 2,209 | 0,000 | Up | AA2 | N |
| **Th0179_002913** | lcl\|JOKZ01000095.1_cds_KKP03872.1_4022 | THAR02_04021 | 2,114 | 0,000 | Up | GH72 | N |
| **Th0179_000189** | lcl\|JOKZ01000014.1_cds_KKP07011.1_891 | THAR02_00890 | 2,105 | 0,001 | Up | GH3 | Y |
| **Th0179_006545** | lcl\|JOKZ01000108.1_cds_KKP03485.1_4406 | THAR02_04405 | 2,087 | 0,013 | Up | CBM1+GH5_5 | Y |
| **Th0179_003586** | lcl\|JOKZ01000257.1_cds_KKP00372.1_7532 | THAR02_07531 | 2,054 | 0,000 | Up | GH20 | Y |
| **Th0179_001277** | lcl\|JOKZ01000064.1_cds_KKP04907.1_3009 | THAR02_03008 | 2,024 | 0,041 | Up | GH55 | Y |
| **Th0179_009090** | lcl\|JOKZ01000270.1_cds_KKP00192.1_7717 | THAR02_07716 | 1,983 | 0,000 | Up | GH64 | N |
| **Th0179_007533** | lcl\|JOKZ01000001.1_cds_KKP07860.1_68 | THAR02_00068 | 1,868 | 0,025 | Up | GH18 | N |
| **Th0179_005496** | lcl\|JOKZ01000072.1_cds_KKP04612.1_3303 | THAR02_03302 | 1,849 | 0,000 | Up | GH16_3 | N |
| **Th0179_011372** | lcl\|JOKZ01000274.1_cds_KKP00125.1_7778 | THAR02_07777 | 1,790 | 0,001 | Up | CBM1+GH18 | Y |
| **Th0179_011548** | lcl\|JOKZ01000410.1_cds_KKO98537.1_9358 | THAR02_09357 | 1,768 | 0,000 | Up | GH2 | N |
| **Th0179_007089** | lcl\|JOKZ01000025.1_cds_KKP06476.1_1435 | THAR02_01434 | 1,749 | 0,000 | Up | GH16_3 | Y |

| | | | | | | | |
|---|---|---|---|---|---|---|---|
| **Th0179_009162** | lcl\|JOKZ01000365.1_cds_KKO99004.1_8898 | THAR02_08897 | 1,736 | 0,000 | Up | CBM1+GH7 | Y |
| **Th0179_005355** | lcl\|JOKZ01000504.1_cds_KKO97791.1_10110 | THAR02_10110 | 1,640 | 0,000 | Up | CE9 | N |
| **Th0179_000330** | lcl\|JOKZ01000530.1_cds_KKO97625.1_10273 | THAR02_10273 | 1,620 | 0,006 | Up | GH17 | N |
| **Th0179_006554** | lcl\|JOKZ01000108.1_cds_KKP03494.1_4415 | THAR02_04414 | 1,525 | 0,000 | Up | CBM1+GH6 | N |

**Supplementary Material 1: Supplementary Table 8.** Expression profiles of CAZymes in the genome of Ta0020 that were differentially expressed under cellulose growth conditions. Transcript expression levels were calculated as TPM values. Genes with p values ≤ 0.05 and fold changes ≥ 1.5 (upregulated) or ≤ -1.5 (downregulated) were considered differentially expressed. Ta0020: *T. atroviride* CBMAI-0020; TaIMI206040: *T. atroviride* IMI206040; FC values: fold change values; DEGs: differentially expressed genes.

| Gene ID_Ta0020 | Protein product_TaIMI206040 | GeneID_TaIMI206040 | FC values | p value | DEG | Family | Secreted |
|---|---|---|---|---|---|---|---|
| **Ta0020_009362** | lcl\|NW_014013636.1_cds_XP_013945973.1_8481 | TRIATDRAFT_154204 | 7,10 | 0,00 | Up | GT1 | N |
| **Ta0020_009111** | lcl\|NW_014013633.1_cds_XP_013943769.1_6607 | TRIATDRAFT_40838 | 6,28 | 0,00 | Up | GT8 | N |
| **Ta0020_009427** | lcl\|NW_014013636.1_cds_XP_013945935.1_8432 | TRIATDRAFT_81098 | 3,00 | 0,01 | Up | CBM42+GH54 | Y |
| **Ta0020_008341** | lcl\|NW_014013635.1_cds_XP_013945238.1_7668 | TRIATDRAFT_217415 | 2,93 | 0,01 | Up | GH18 | Y |
| **Ta0020_009145** | lcl\|NW_014013633.1_cds_XP_013943809.1_6645 | TRIATDRAFT_41194 | 2,78 | 0,00 | Up | GH64 | Y |
| **Ta0020_005527** | lcl\|NW_014013631.1_cds_XP_013942515.1_4982 | TRIATDRAFT_150889 | 2,62 | 0,00 | Up | GH5_9 | N |
| **Ta0020_009301** | lcl\|NW_014013633.1_cds_XP_013943791.1_6811 | TRIATDRAFT_41039 | 2,58 | 0,00 | Up | GH20 | Y |
| **Ta0020_004225** | lcl\|NW_014013632.1_cds_XP_013942791.1_5511 | TRIATDRAFT_16857 | 2,20 | 0,00 | Up | GH75 | Y |
| **Ta0020_004226** | lcl\|NW_014013632.1_cds_XP_013942791.1_5511 | TRIATDRAFT_16857 | 2,20 | 0,00 | Up | GH75 | N |
| **Ta0020_005731** | lcl\|NW_014013631.1_cds_XP_013942693.1_5206 | TRIATDRAFT_285140 | 1,99 | 0,00 | Up | GH76 | Y |
| **Ta0020_004068** | lcl\|NW_014013632.1_cds_XP_013942958.1_5347 | TRIATDRAFT_88379 | 1,95 | 0,00 | Up | AA2 | Y |
| **Ta0020_006050** | lcl\|NW_014013637.1_cds_XP_013946279.1_8765 | TRIATDRAFT_81867 | 1,82 | 0,01 | Up | GH5_24 | N |
| **Ta0020_001760** | lcl\|NW_014013630.1_cds_XP_013940821.1_3151 | TRIATDRAFT_161159 | 1,77 | 0,03 | Up | GH3 | Y |

| | | | | | | | |
|---|---|---|---|---|---|---|---|
| **Ta0020_004278** | lcl\|NW_014013632.1_cds_XP_013942813.1_5569 | TRIATDRAFT_37969 | 1,66 | 0,00 | Up | GH16_3 | Y |

**Supplementary Material 1: Supplementary Table 9.** Expression profiles of CAZymes in the genome of Tr0711 that were differentially expressed under cellulose growth conditions. Transcript expression levels were calculated as TPM values. Genes with p values ≤ 0.05 and fold changes ≥ 1.5 (upregulated) or ≤ -1.5 (downregulated) were considered differentially expressed. Tr0711: *T. reesei* CBMAI-0711; Tr2.0: *T. reesei* v2.0; FC values: fold change values; DEGs: differentially expressed genes.

| Gene ID_Tr0711 | GeneID_Trv2.0 | FC values | p value | DEG | Family | Secreted |
|---|---|---|---|---|---|---|
| **Tr0711_006897** | jgi\|Trire\|80340 | 14,68 | 0,00 | Up | GT32 | N |
| **Tr0711_007407** | jgi\|Trire\|108776 | 7,26 | 0,00 | Up | GH55 | Y |
| **Tr0711_008576** | jgi\|Trire\|108776 | 7,26 | 0,00 | Up | GH55 | N |
| **Tr0711_000934** | jgi\|Trire\|106575 | 6,99 | 0,00 | Up | GH79 | Y |
| **Tr0711_004222** | jgi\|Trire\|119859 | 6,72 | 0,00 | Up | GH18 | Y |
| **Tr0711_004484** | jgi\|Trire\|49976 | 5,38 | 0,00 | Up | CBM1+GH45 | Y |
| **Tr0711_000520** | jgi\|Trire\|38441 | 5,11 | 0,00 | Up | GH132 | Y |
| **Tr0711_007923** | jgi\|Trire\|104664 | 4,64 | 0,00 | Up | GH132 | Y |
| **Tr0711_002287** | jgi\|Trire\|64375 | 3,23 | 0,02 | Up | GH5_9 | N |
| **Tr0711_000101** | jgi\|Trire\|76700 | 2,47 | 0,03 | Up | GH17 | N |
| **Tr0711_000742** | jgi\|Trire\|103850 | 2,42 | 0,03 | Up | GT33 | N |
| **Tr0711_006819** | jgi\|Trire\|70845 | 2,20 | 0,01 | Up | GH55 | Y |
| **Tr0711_008543** | jgi\|Trire\|71563 | 2,08 | 0,00 | Up | GT2 | N |
| **Tr0711_002443** | jgi\|Trire\|73638 | 1,79 | 0,01 | Up | CBM1 | Y |

**Supplementary Material 1: Supplementary Table 10.** Differentially expressed CAZyme-encoding genes under cellulose growth conditions and their orthologs in all evaluated strains. Transcript expression levels were calculated as TPM values. The orthologs of the CAZyme-encoding genes among Th3844, Th0179, Ta0020, and Tr0711 were identified from the OrthoFinder results. Genes with p values ≤ 0.05 and fold changes ≥ 1.5 (upregulated) or ≤ -1.5 (downregulated) were considered differentially expressed. Th3844: *T. harzianum* IOC-3844; Th0179: *T. harzianum* CBMAI-0179; Ta0020: *T. atroviride* CBMAI-0020; Tr0711: *T. reesei* CBMAI-0711; FC values: fold change values.

| | Gene ID | | | | | | | |
|---|---|---|---|---|---|---|---|---|
| **CAZyme family** | **Th3844** | **Th0179** | **Ta0020** | **Tr0711** | **FC values Th3844** | **FC values Th0179** | **FC values Ta0020** | **FC values Tr0711** |
| **GH11** | 001491 | 010414 | 000032 | - | 3,30 | 1,36 | -5,13 | - |
| **AA9+CBM1** | 001506 | 010399 | 000021 | - | 5,23 | 2,60 | -4,97 | - |
| **GH72** | 003274 | 002913 | 003744 | - | 1,95 | 2,11 | -1,63 | - |
| **GH1** | 003759 | 003380 | 003310 | - | 1,79 | 2,64 | -1,68 | - |
| **GH20** | 003967 | 003586 | 003822 | - | -1,28 | 2,05 | -1,44 | - |
| **GT4** | 005396 | 004687 | 007475 | - | -1,35 | -1,43 | 1,48 | - |
| **AA11** | 005513 | 004573 | 007354 | - | 1,52 | -1,71 | -2,05 | - |
| **GH16_3** | 006635 | 005496 | 005640 | - | 1,83 | 1,85 | -1,44 | - |
| **GH1** | 007252 | 004888 | 005032 | - | 1,93 | 2,22 | -2,23 | - |
| **GH106+GH10** | 008264 | 009135 | 004152 | - | 5,30 | 2,25 | -6,95 | - |
| **CBM1+GH7** | 008290 | 009162 | 009328 | - | 3,42 | 1,74 | -3,79 | - |
| **GH20** | 008322 | 009190 | 009301 | - | -1,23 | -1,56 | 2,58 | - |
| **GH64** | 009030 | 009090 | 008705 | - | -1,15 | 1,98 | 1,43 | - |
| **CBM1+GH6** | 010327 | 006554 | 007547 | - | 3,92 | 1,52 | -3,30 | - |
| **AA3_2** | 011000 | 011191 | 000129 | 006613 | -1,60 | -2,53 | -2,29 | -5,58 |

| | Th3844 | Th0179 | Ta0020 | Tr0711 |
|---|---|---|---|---|
| GH134 | 3 | 0 | 0 | 0 |
| GH49 | 3 | 0 | 0 | 0 |
| PL38 | 0 | 2 | 1 | 1 |
| GH17 | 1 | 1 | 2 | 1 |
| GH63 | 1 | 1 | 2 | 1 |
| GH67 | 0 | 2 | 2 | 1 |
| GH89 | 1 | 1 | 1 | 2 |
| AA4 | 2 | 3 | 0 | 0 |
| GH10 | 2 | 2 | 1 | 1 |
| GH154 | 1 | 2 | 2 | 1 |
| GH38 | 2 | 2 | 1 | 1 |
| GH62 | 1 | 2 | 2 | 1 |
| GH65 | 1 | 1 | 2 | 2 |
| GH78 | 2 | 2 | 2 | 1 |
| GH88 | 2 | 3 | 2 | 0 |
| AA7 | 2 | 3 | 1 | 1 |
| GH132 | 2 | 2 | 2 | 2 |
| GH37 | 2 | 2 | 2 | 2 |
| GH54 | 2 | 2 | 2 | 2 |
| GH7 | 2 | 2 | 2 | 2 |
| GH81 | 2 | 2 | 2 | 2 |
| GT34 | 2 | 2 | 2 | 2 |
| GT57 | 2 | 2 | 2 | 2 |
| GT64 | 2 | 2 | 2 | 2 |
| PL20 | 2 | 2 | 2 | 2 |
| CE16 | 2 | 2 | 2 | 2 |
| AA14 | 2 | 2 | 2 | 2 |
| GH125 | 3 | 2 | 2 | 2 |
| GH15 | 2 | 2 | 3 | 2 |
| GH79 | 2 | 2 | 2 | 3 |
| GH12 | 3 | 3 | 2 | 2 |
| GT15 | 2 | 3 | 3 | 3 |
| GT62 | 3 | 3 | 2 | 3 |
| PL7 | 5 | 3 | 1 | 2 |
| GH20 | 3 | 3 | 3 | 3 |
| GT39 | 3 | 3 | 3 | 3 |
| GT8 | 3 | 3 | 3 | 3 |
| CE4 | 4 | 3 | 3 | 2 |
| AA2 | 1 | 3 | 5 | 3 |
| AA9 | 3 | 3 | 3 | 3 |
| GH128 | 2 | 4 | 4 | 3 |
| AA11 | 5 | 3 | 2 | 3 |
| GH1 | 4 | 4 | 4 | 2 |
| GH11 | 4 | 4 | 4 | 3 |
| GH43 | 5 | 4 | 4 | 2 |
| CE3 | 4 | 4 | 3 | 4 |
| CE5 | 4 | 4 | 3 | 4 |
| GT22 | 4 | 4 | 4 | 4 |
| GT69 | 4 | 4 | 4 | 4 |
| GH64 | 8 | 3 | 3 | 3 |
| GH95 | 4 | 5 | 4 | 4 |
| GT20 | 5 | 4 | 4 | 4 |
| GH72 | 3 | 5 | 5 | 5 |
| GH28 | 5 | 5 | 5 | 4 |
| GH75 | 5 | 5 | 6 | 3 |
| GT32 | 4 | 5 | 5 | 5 |
| GT4 | 4 | 5 | 5 | 5 |
| GH27 | 7 | 9 | 3 | 1 |
| GH30 | 4 | 6 | 5 | 5 |
| GH31 | 4 | 5 | 7 | 4 |
| GT1 | 7 | 5 | 5 | 4 |
| GH71 | 8 | 6 | 4 | 4 |
| GT90 | 6 | 6 | 7 | 6 |
| GH13 | 9 | 7 | 5 | 5 |
| GH92 | 7 | 7 | 8 | 6 |
| AA1 | 9 | 9 | 6 | 4 |
| GH47 | 7 | 8 | 8 | 8 |
| GH55 | 11 | 10 | 8 | 6 |
| GH5 | 10 | 10 | 9 | 7 |
| GH76 | 11 | 9 | 9 | 8 |
| GH2 | 11 | 13 | 10 | 7 |
| GT2 | 11 | 10 | 13 | 13 |
| GH16 | 13 | 13 | 13 | 13 |
| GH3 | 16 | 17 | 15 | 12 |
| AA3 | 19 | 20 | 12 | 12 |
| GH18 | 29 | 26 | 25 | 18 |

**Supplementary Material 1: Supplementary Figure 5. Quantitative comparison of the CAZyme repertoires of *Trichoderma* isolates.** Heatmap of the number of enzymes of each CAZY family of Th3844, Th0179, Ta0020, and Tr0711. This map includes all the enzymes/proteins present in the evaluated genomes. Ta0020: *T. atroviride* CBMAI-0020; Th0179: *T. harzianum* CBMAI-0179; Th3844: *T. harzianum* IOC-3844; Tr0711: *T. reesei* CBMAI-0711.

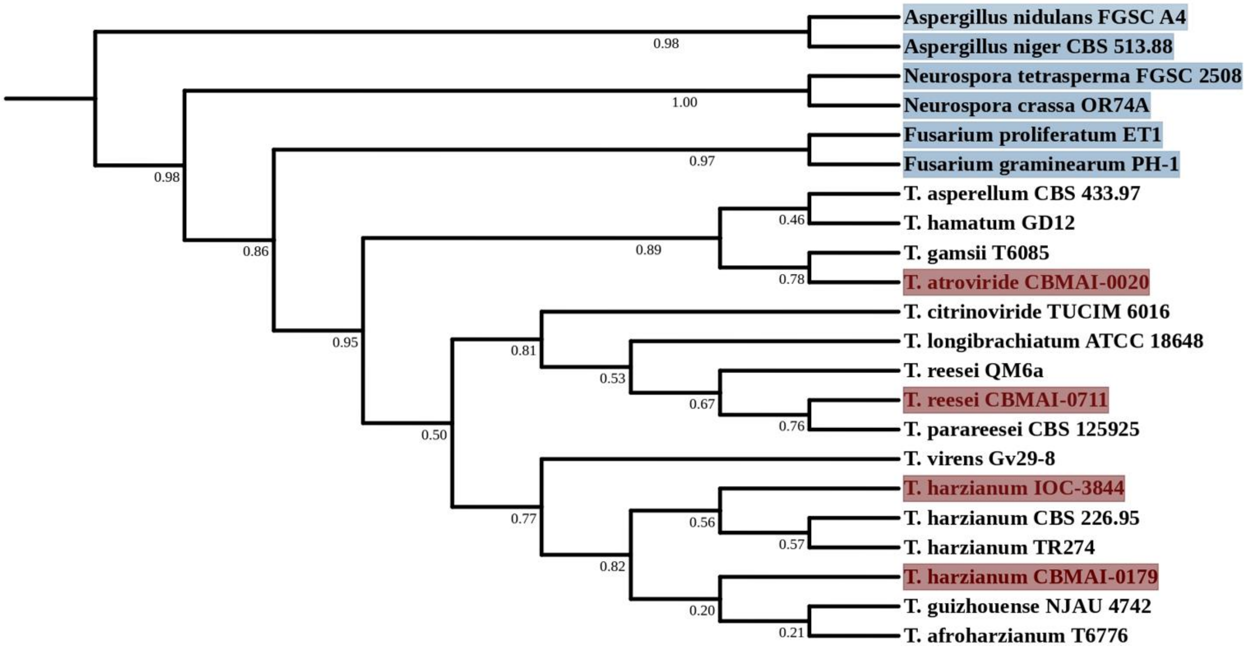


**Supplementary Material 1: Supplementary Figure 6. Phylogenetic relationships of *Trichoderma* spp. as determined by orthology analysis.** The phylogenetic tree modeled by OrthoFinder software was based on the concatenation of 2,195 single-copy orthogroups. This methodology illustrated the inferred relationships among 18 *Trichoderma* spp. whose proteomes are available in the NCBI database. *Fusarium* spp., *Aspergillus* spp., and *Neurospora* spp. were used as outgroups. Bootstrap values are shown at the nodes.

**Gene markers used for the identification of the evaluated strains and their respective nucleotide sequences**

***>Th3844_tef1***

AAGAACATGATCACTGGTACTTCCCAGGCCGATTGCGCTATCCTCATCATTGCCGCC
GGTACTGGTGAGTTCGAGGCTGGTATCTCCAAGGATGGCCAGACCCGTGAGCACGC
TCTGCTCGCCTACACCCTGGGTGTCAAGCAGCTCATTGTTGCCATCAACAAGATGGA
CACTGCCAACTGGGCCGAGGCTCGTTACCAGGAAATCATCAAGGAGACTTCCAACT
TCATCAAGAAGGTCGGCTTCAACCCCAAGGCTGTTGCTTTCGTCCCCATCTCCGGCT
TCAACGGTGACAACATGCTCCAGCCCTCCACCAACTGCCCCTGGTACAAGGGCTGG
GAGAAGGAGACCAAGGCTGGCAAGTTCACCGGCAAGACCCTCCTTGAGGCTATCGA
CTCCATCGAGCCCCCCAAGCGTCCCACGGACAAGCCCCTCCGTCTTCCCCTCCAGGA
TGTCTACAAGATCGGTGGTATTGGAACAGTTCCCGTCGGCCGTATCGAGACTGGTAT
CCTCAAGCCCGGTATGGTCGTCACCTTCGCTCCCTCCAACGTCACCACTGAAGTCAA
GTCCGTCGAGATGCACCACGAGCAGCTCGTCGAGGGTGTTCCCGGTGACAA

***>Th0179_tef1***

AACATGATCACTGGTACTTCCCAGGCTGACTGCGCCATTCTCATCATTGCCGCCGGT
ACTGGTGAGTTCGAGGCTGGTATCTCCAAGGATGGCCAGACTCGTGAGCACGCTCTG
CTCGCCTACACCCTGGGTGTCAAGCAGCTCATCGTTGCCATCAACAAGATGGACACT
GCCAACTGGGCCGAGGCTCGTTACCAGGAAATCATCAAGGAGACTTCCAACTTCAT
CAAGAAGGTCGGCTTCAACCCCAAGGCTGTTGCTTTCGTCCCCATCTCCGGTTTCAA
CGGTGACAACATGCTCCAGCCCTCCACCAACTGCCCCTGGTACAAGGGTTGGGAGA

AGGAGACCAAGGCTGGCAAGTTCACCGGCAAGACCCTCCTTGAGGCCATCGACTCC

ATCGAGCCCCCCAAGCGTCCCACGGACAAGCCCCTCCGTCTTCCCCTCCAGGATGTC

TACAAGATCGGTGGTATCGGAACAGTTCCCGTCGGCCGTATCGAGACTGGTGTCCTC

AAGCCCGGTATGGTCGTCACCTTCGCTCCCTCCAACGTCACCACTGAAGTCAAGTCC

GTCGAGATGCACCACGAGCAGCTCACCGAGGGTGTTCCCGGTGACAA

***>Tr0711_tef1***

GTTGTGCCGACAGGTTTTTTTTCATCACCCCGCTTTCTTCTACCCCTCCGAGCGACgC

AAATTTTTTTGCTGCCTTACGATGGGTTTTAGTGGG

***>Ta0020_tef1***

CAAGAACATGATCACTGGTACCTCCCAGGCTGACTGCGCTATCCTGATTATCGCTGC

CGGTACTGGTGAGTTCGAGGCTGGTATCTCCAAGGATGGCCAGACCCGTGAGCACG

CTCTGCTCGCCTACACCCTGGGTGTCAAGCAGCTCATTGTTGCCATCAACAAGATGG

ACACTGCCAACTGGGCCGAGGCTCGTTACCTTGAGATCATCAAGGAGACCTCCAACT

TCATCAAGAAGGTCGGCTTCAACCCCAAGACCGTTGCCTTCGTCCCCATCTCCGGCT

TCAACGGCGACAACATGTTGACTGCCTCCACCAACTGCCCCTGGTACAAGGGCTGG

GAGAAGGAGACCAAGGCTGGCAAGTCCACCGGCAAGACCCTTCTCGAGGCCATTGA

CGCCATTGAGCCCCCCAAGCGTCCCACAGACAAGCCCCTCCGTCTTCCCCTTCAGGA

TGTTTACAAGATCGGTGGTATCGGAACTGTCCCTGTCGGCCGTATCGAGACTGGTGT

CCTCAAGCCCGGTATGGTCGTTACCTTTGCTCCCTCCAACGTGACCACTGAAGTCAA

GTCCGTTGAGATGCACCACGAGCAGCTTACCGAGGGTGTCCCCGGTGACAA

***>Th3844_006646-RA_rpb2***

ATGGCTGACTACGAGGACGAATACGACTACGAGAACTACGGAGACGAGGATGAAG

GCATCACTCCCGAGGATTGCTGGACTGTCATCTCCTCCTTCTTCGAAACCAAGGGTC

TCGTATCGCAGCAGACCGACTCTTTTGACGAATTCACCCAGACGACGATCCAGGACC

TCGTAAACGAATACTCCACCATCACACTCGACCAGCCAAACCCTCCTTCGCCACCCG

GCCGAACAATAGCCCTTCGTCGATATGAGATCAAGTTTGGAAGCGTCATGGTGTCAC

GTCCCACTATCAGTGAGACGGACGGCACTGTGACTTCTCTGCTCCCTTACGAGTGCC

GAGACCGTAACTTGACCTACGCCAGTCCCCTCTATATCAAGATCACTAAAAAGGTGT

CGGCTGCTGTTGAGAGGGAGGTTCCCCTGCACGAAATGGACGATGCTCAGCAGGAG

GAATACGCAAGAACCGGCGAACACCCTACAAAGCTCGAGTGGGAAGAGGAGGAGA

ACGGCGAAGATGACAACATCGGCAAGTCTGATGACTGGAAGGACATGGTCTTCGTT

GGCAAGCTGCCCATCATGGTCAAATCCAAGATTTGTCATCTGAGCCGTGAACAGGAT

GATAGCCTGTTCCTTGTCAACGAATGTCCCTACGATCAGGGTGGCTACTTTGTTATCA

ACGGCAGTGAAAAGGTCCTCATCGCCCAAGAGAGATCCGCCGCCAACATTGTCCAA

GTCTTCAAGAAGGCCCAGCCCAGTGCCTATACCTATACGGCTGAAATCCGAAGTGC

GCTGGAAAAGGGCTCACGGCTCATCTCTAGCATGATGCTCAAGCTGTATGGCAAGG

GAGACTCTGCTCGTGGTGGCTTTGGCCAGACTATCCACACCACCCTGCCCTTTGTCA

AGTCAGATCTTCCCGTCGCCATTGTCTTCCGTGCCCTGGGTGTCGTTTCTGATGAAGA

TATCCTCAACCACATTTGCTACGACCGCAACGACAGCCAGATGCTGGAGATGCTTCG

ACCTTGTATTGAGGAGGCCTTCTGTGTCCAGGACCGAGAGGTTGCTCTGGATTTCAT

CGGAAAGCGTGGAAACCGAGACCAAGCTGGTCTCGGACGCGAGAAGCGTGTCCGCG

TGGCCAAGGATATCCTTCAGAAGGAGACTCTTCCCCACATTTCACAGACAGAGGGA

AGTGAAACCAGAAAGGCATTTTTCTTGGGATACATGGTGCACAAGCTGTTGCAATGT

GCGCTCGGAAGAAGAGAGCCCGACGATCGTGACCACTTTGGAAAGAAGCGTCTGGA
TCTGGCGGGTCCCCTGCTGGCAAAGCTGTTCCGTGGTATCATGCGAAGGATGAACAC
TGAGTTGGCCAACTATCTGAGACGATGCGTCGAGGGCAACCGACACTTCAACCTTGC
TGTGGGTATCAAGCCCGGCACGCTTTCAAACGGATTGAAGTATTCGCTTGCCACAGG
AAACTGGGGTGATCAGAAGAAGGCCATGAGCTCAACTGCAGGTGTGTCCCAGGTGC
TTAACCGTTACACGTTTGCTTCGACCCTATCACATTTGCGTCGTACCAATACTCCTAT
CGGAAGAGATGGTAAGCTCGCAAAGCCTCGACAGCTTCACAACACGCACTGGGGTT
TGGTCTGCCCGGCCGAGACACCCGAGGGACAGGCTTGTGGTCTGGTCAAGAACTTGT
CTTTGATGTGTTACGTCAGTGTCGGTTCTCCCTCCGAACCTCTGATTGAGTTCATGAT
CAACAGAGGTATGGAAGTCGTGGAAGAGTACGAGCCGCTGCGGTATCCTCATGCTA
CAAAGATTTTTGTGAACGGTGTCTGGGTTGGAGTCCACCAAGACCCTAAGCACTTGG
TGAACCAGGTCCTGGACACTCGTCGCAAGTCCTATCTGCAATACGAAGTCTCTCTCG
TGAGAGAAATTCGAGACCAGGAATTCAAAATCTTTTCCGACGCAGGCCGTGTAATG
CGGCCAGTCTTTACCGTTCAGCAGGAAGATGACCCGGAAACGGGCATCAACAAGGG
CCACCTGGTATTGACCAAGGAGCTCGTCAATAGATTGGCCAAGGAGCAGGCTGAAC
CTCCGGAAGACCCCAGCATGAAGATTGGATGGGAGGGATTGATTAGGGCTGGTGCG
GTTGAATATCTCGACGCCGAGGAAGAGGAGACGTCCATGATCTGCATGACGCCAGA
GGATCTCGAGCTGTATCGTCTTCAGAAGGCTGGTATTAACACTGAGGAAGACATGG
GAGATGACCCGAACAAGCGACTAAAGACCAAGACAAACCCGACTACTCACATGTAC
ACCCATTGCGAGATTCACCCAAGTATGATCTTAGGCATCTGTGCTAGTATCATTCCTT
TCCCCGATCACAACCAGTCCCCCCGTAACACTTACCAATCTGCCATGGGTAAGCAAG
CTATGGGTTTCTTCCTCACAAACTATTCCCGGCGCATGGACACCATGGCCAACATTC
TCTACTACCCCCAGAAGCCGCTGGGTACCACTCGATCCATGGAGTTTTTGAAGTTCC

GAGAGTTGCCTGCTGGTCAGAACGCCATTGTAGCCATTGCCTGTTACTCTGGTTACA

ACCAGGAAGATTCCGTCATTATGAACCAGAGTAGTATTGACAGAGGTCTTTTCCGAA

GTCTCTTCTTCCGATCATACTCGGATCAGGAGAAGAAGGTTGGATTGAACTACACCG

AAGTGTTTGAGAAGCCATTCCACCAAAACACTCTTCGCATGAAGCACGGCACATAC

GACAAGCTGGATGAAGACGGTATCGTTGCTCCTGGTGTCCGAGTATCTGGTGAAGAT

ATCATTATTGGCAAGACGGCGCCGATTGACGCTGAGACACAAGATCTCGGTACCAG

GACGACTATGCACCAGAGGCGAGATATCTCAACTCCTCTGCGTAGCACTGAGAACG

GTATCGTTGACTCTGTCATTGTGACAGTGAACGCGGACAATGTCAAGTATGTCAAGG

TCCGTGTCCGAACGACCAAGATTCCTCAGATTGGAGACAAGTTCGCCTCTCGTCACG

GTCAAAAGGGCACTATTGGTGTTACTTACCGACAAGAGGACATGCCCTTCACAAGA

GAAGGTGTCACTCCAGACATCATCATCAACCCTCACGCCATTCCATCCCGTATGACA

ATCGCTCACTTGATTGAGTGTCTCCTAAGTAAGGTCTCTACGCTGGAAGGTATGGAG

GGTGATGCCACCCCCTTCACGGATGTCACCGTCGACTCGGTCTCAGAGCTGTTGCGA

AAGCACGGCTACCAGTCACGAGGCTTTGAGATTATGTACAATGGCCACACAGGCCG

CAAGCTGAGAGCCCAGGTGTTCTTTGGACCAACATACTACCAGCGACTCCGTCACAT

GGTGGACGACAAGATCCACGCTCGTGCCCGTGGTCCCGTGCAGATTATGACAAGAC

AACCCGTGGAGGGTCGTGCCAGAGATGGTGGTCTCCGATTCGGAGAAATGGAACGT

GATTGTATGATTGCTCACGGTGCCGCGTCCTTCCTCAAGGAGCGATTGTTTGAGGTG

TCAGACGCCTTCCGAGTTCACATTTGCGAGATTTGTGGACTCATGACGCCTATTGCC

AACCTCTCTAAACAATCGTTCGAGTGTCGACCTTGTAAGAACAAGACCAAGATTGCA

CAGATTCACATCCCTTATGCCGCCAAGCTCCTGTTCCAGGAACTCCAGTCGATGAAC

ATTGCGGCTCGCATGTTCACAAACCGGTCTGGCGCGTCCATCAGGTAA

***>Th0179_005485-RA_rpb2***

ATGGCCGACTACGAGGACGACTACGACTACGAGAACTACGGAGACGAGGATGAAG

GCATCACTCCCGAGGATTGCTGGACCGTCATCTCCTCCTTCTTCGAAACCAAGGGTC

TTGTATCGCAGCAGACAGACTCCTTTGACGAATTCACGCAGACGACCATCCAGGATC

TCGTAAACGAATACTCCACCATCACACTCGACCAGCCAAACCCTCCCTCGCCACCCG

GCCGAACAATAGCCCTTCGTCGATATGAGATCAAGTTTGGAAGCGTCATGGTGTCAC

GTCCCACCATCAGTGAGACGGACGGAACCGTGACTTCTCTGCTCCCTTACGAGTGCC

GAGACCGCAACTTGACGTACGCCAGTCCCCTCTACATCAAGATTACTAAAAAGGTGT

CGGCTGCTGTTGAGAGGGAGGTTCCCCTGCACGAAATGGACGATGCCCAGCAGGAA

GAATACGCAAGAACCGGCGAACACCCTACAAAGCTTGAGTGGGAAGAGGAGGAGA

ACGGCGAGGATGACAACATCGGCAAGTCTGATGATTGGAAGGACATGGTCTTCGTT

GGCAAGCTGCCCATCATGGTCAAATCCAAGATTTGTCATCTGAGCCGTGAACAGGAT

GATAGCCTGTTCCTTGTCAACGAATGTCCCTACGATCAGGGTGGCTACTTTGTTATCA

ACGGCAGTGAAAAGGTTCTCATCGCCCAAGAGAGATCTGCCGCCAACATTGTCCAA

GTCTTCAAGAAGGCCCAGCCCAGTGCCTATACCTATACGGCTGAAATCCGAAGTGC

GTTGGAAAAGGGATCACGGCTCATCTCCAGCATGATGCTCAAGCTGTATGGCAAGG

GAGACTCTGCTCGTGGTGGCTTTGGCCAGACTATCCACACCACTCTACCCTTTGTCA

AGTCAGATCTTCCCGTCGCCATTGTCTTCCGTGCCCTGGGTGTCGTTTCTGATGAAGA

TATCCTCAACCACATTTGCTACGACCGCAACGACAGCCAGATGCTGGAGATGCTTCG

ACCTTGTATTGAGGAGGCCTTCTGTGTCCAGGACCGAGAGGTTGCTCTCGATTTCAT

CGGAAAGCGTGGAAACCGAGACCAAGCTGGTCTCGGACGTGAGAAGCGTGTCCGCG

TGGCCAAGGATATCCTTCAGAAGGAGACTCTTCCCCACATTTCGCAGACAGAGGGA

AGTGAAACCAGAAAGGCATTCTTCCTGGGATACATGGTGCACAAGCTGTTGCAGTGT

GCGCTCGGAAGAAGAGAGCCCGACGATCGTGACCACTTTGGAAAGAAGCGTCTGGA

TCTGGCGGGTCCCCTGCTGGCCAAGCTGTTCCGTGGTATCATGCGAAGGATGAATAC

TGAGTTGGCCAACTACCTGAGACGATGCGTTGAGGGCAACCGACACTTCAACCTTGC

TGTTGGTATCAAGCCCGGCACGCTTTCAAACGGATTGAAGTATTCGCTTGCCACAGG

CAACTGGGGTGATCAGAAGAAGGCCATGAGCTCAACTGCAGGTGTGTCCCAGGTGC

TTAACCGATACACGTTTGCTTCGACCCTGTCACATTTGCGTCGTACCAACACTCCCAT

CGGAAGAGATGGTAAGCTGGCGAAGCCTCGACAGCTTCACAACACGCATTGGGGTT

TGGTCTGCCCAGCCGAGACACCCGAAGGACAGGCCTGTGGTCTGGTCAAGAACCTG

TCTTTGATGTGTTACGTCAGTGTCGGTTCTCCCTCCGAGCCTCTGATTGAGTTCATGA

TCAACAGAGGTATGGAAGTCGTCGAAGAGTATGAGCCTCTGCGGTATCCTCATGCTA

CAAAGATTTTTGTGAACGGTGTCTGGGTTGGAGTTCACCAAGACCCTAAGCACTTGG

TGAACCAGGTTCTAGATACTCGTCGCAAGTCCTATCTGCAATACGAAGTCTCTCTCG

TGAGAGAAATTCGAGACCAGGAATTCAAAATCTTTTCCGACGCAGGTCGTGTCATGC

GACCAGTCTTTACCGTTCAGCAGGAAGATGACCCGGAAACGGGCATCAACAAGGGC

CACCTGGTATTGACCAAGGAGCTCGTCAATAGATTGGCTAAGGAGCAGGCTGAGCC

TCCGGAAGACCCCAGCATGAAGATTGGATGGGAGGGATTGATTAGGGCTGGTGCAG

TTGAATATCTCGACGCTGAGGAAGAGGAGACGTCCATGATCTGCATGACGCCAGAG

GATCTCGAGCTGTATCGTCTTCAGAAGGCCGGTATTAACACTGAGGAAGACATGGG

AGATGACCCGAACAAGCGACTGAAGACCAAGACGAACCCGACAACTCACATGTACA

CCCATTGCGAGATTCACCCAAGTATGATCTTAGGTATCTGTGCTAGTATCATTCCTTT

CCCCGATCACAACCAGTCTCCCCGTAACACTTACCAATCTGCCATGGGTAAGCAAGC

TATGGGTTTCTTCCTCACGAACTATTCCCGGCGCATGGACACCATGGCCAACATCCT

TTACTACCCTCAGAAGCCGCTGGGTACCACTCGATCCATGGAGTTTTTGAAATTCCG

AGAGTTACCTGCTGGTCAGAACGCCATTGTAGCCATTGCCTGCTACTCTGGCTACAA

CCAGGAAGATTCCGTCATTATGAACCAGAGTAGTATTGACAGAGGTCTGTTCCGAAG

TCTCTTCTTCCGATCATACTCGGATCAGGAGAAGAAGGTTGGATTGAACTACACCGA

AGTGTTTGAGAAGCCCTTCCACCAAAACACTCTTCGCATGAAGCACGGTACATATGA

CAAGCTGGATGAAGACGGTATCGTTGCTCCTGGTGTCCGAGTATCTGGTGAAGACAT

CATTATTGGCAAGACGGCGCCGATTGATGCTGAGACACAAGATCTCGGCACCAGGA

CGACTATGCACCAGAGGCGAGATATCTCAACTCCTCTGCGTAGCACAGAGAACGGT

ATCGTTGACTCTGTCATCGTGACAGTGAACGCGGACAATGTCAAGTATGTCAAGGTC

CGTGTCCGAACGACCAAGATTCCCCAGATTGGAGACAAGTTCGCCTCTCGTCACGGT

CAAAAGGGCACTATTGGTGTTACTTACCGACAAGAGGACATGCCCTTCACAAGAGA

AGGTGTCACTCCAGACATCATCATCAACCCTCACGCCATTCCGTCCCGTATGACAAT

CGCTCACTTGATTGAGTGTCTCCTAAGTAAGGTCTCTACGCTGGAAGGTATGGAGGG

TGATGCCACTCCCTTTACGGATGTCACGGTCGACTCGGTCTCAGAGCTGTTGCGAAA

GCACGGTTACCAGTCACGAGGCTTTGAGATTATGTATAATGGCCATACAGGCCGCAA

GCTGAGAGCCCAGGTGTTCTTCGGACCAACATACTACCAGCGACTCCGTCACATGGT

GGACGACAAGATCCACGCTCGTGCCCGTGGTCCCGTGCAGATCATGACTAGACAAC

CTGTGGAAGGTCGTGCGAGAGATGGTGGTCTCCGATTCGGAGAAATGGAACGTGAT

TGTATGATTGCTCACGGTGCCGCGTCCTTCCTCAAGGAGCGATTGTTTGAGGTGTCA

GACGCCTTCCGAGTTCACATCTGCGAGATTTGTGGACTCATGACGCCTATTGCCAAC

CTTTCTAAGCAATCGTTCGAGTGTCGACCTTGTAAGAACAAGACCAAGATTGCGCAG

ATTCACATCCCTTATGCTGCCAAGCTCCTGTTCCAGGAACTCCAGTCCATGAACATT

GCGGCTCGCATGTTCACAAACCGGTCTGGCGCGTCTATCAGGTAA

***>Tr0711_007464-RA_rpb2***

ATGGCTGATTACGAGGACGACTACGACTACGAAAACTATGGAGACGAGGACGAGGG

CATCACTCCTGAGGACTGCTGGACCGTCATCTCCTCCTTCTTCGAGACCAAGGGCCT

TGTCTCGCAGCAGACAGACTCCTTTGACGAGTTCACGCAGACGACAATCCAGGACCT

CGTAAACGAATACTCCACCATCACCCTCGACCAGCCCAACCCGCCTTCGCCGCCCGG

CCGGACAATAGCCCTTCGTCGATATGAGATCAAGTTTGGAAGCGTCATGGTGTCACG

CCCCACTATCAGTGAGACGGACGGAACCGTCACCTCCCTGCTTCCGTACGAGTGCCG

AGATCGCAACTTGACATACGCCAGTCCGCTTTACATTAAGATTACCAAGAAGCTGTC

GGCCGCCGTCGAGAAGGAGATTCCGTTGCACGAGATGGACGACGCCCAGCAGGAGG

AGTACGCAAGGACCGGCGAGGCCCCGACAAAACTTGAGTGGGAGGAGGAGGAGGC

TGGCGAAGATGACCACAACATTGGCAAGTCTGAAGACTGGAAGGATATGGTTTTCG

TGGGCAAGCTGCCCATCATGGTCAAGTCCAAGATCTGTCATTTGAGCCGCGAGACCG

ACGACAGCCTGTTTCTCGTCAACGAGTGTCCCTACGACCAGGGCGGCTACTTTGTCA

TCAATGGCAGTGAAAAAGTTCTCATCGCCCAGGAGAGATCTGCCGCCAACATCGTTC

AAGTGTTTAAAAAGGCCCAGCCTAGTGCCTACACCTACACGGCGGAGATTCGAAGT

GCGCTCGAGAAGGGCTCGAGGCTCATCTCTAGCATGATGCTCAAGCTCTACGGCAA

GGGAGATTCTGCCCGCGGAGGCTTTGGTCAGACCATTCACACGACCCTGCCCTTTGT

CAAGTCGGATCTTCCGGTAGCCATTGTCTTCCGCGCCTTGGGTGTCGTTTCTGATGAA

GACATCCTGAACCATATTTGCTACGACCGAAACGACAGTCAGATGCTGGAAATGCTT

CGGCCATGCATTGAGGAGGCGTTCTGTGTCCAGGACCGAGAGGTCGCTCTGGACTTC

ATCGGAAAGCGTGGTAACCGAGATCAAGCCGGTCTCGGCCGTGAGAAGCGTGTCCG

TGTGGCCAAGGATATCCTCCAAAAGGAGACGCTGCCTCACATTTCCCAGACAGAAG

GTAGCGAGACCAGAAAGGCCTTCTTCTTGGGCTACATGGTGCACAAACTGCTGCAGT

GCGCACTCGGCAGAAGAGAACCCGACGATCGTGACCACTTTGGAAAGAAGCGGCTG
GACCTGGCGGGTCCTCTGCTGGCCAAGCTGTTCCGTGGCATCATGCGAAGAATGAAC
ACCGAGCTGGCCAACTATCTGAGACGGTGCGTGGAGGGCAACCGACACTTCAATCT
CGCCGTCGGCATCAAGCCCGGCACGCTTTCAAACGGCCTAAAGTACTCGCTCGCCAC
TGGAAACTGGGGTGATCAGAAGAAGGCCATGAGCTCGACCGCAGGTGTGTCTCAGG
TGCTCAACCGCTACACGTTTGCCTCGACCCTCTCGCATTTGCGGCGTACCAACACGC
CTATCGGAAGAGATGGCAAGCTGGCGAAGCCTCGACAGCTTCACAACACCCACTGG
GGTCTCGTCTGCCCGGCCGAGACACCCGAAGGACAGGCCTGTGGTCTTGTCAAGAA
CCTGTCTCTGATGTGTTATGTCAGTGTCGGCTCTCCGTCAGAGCCGTTGATTGAGTTT
ATGATCAATAGGGGCATGGAAGTGGTCGAGGAATACGAGCCACTGCGTTATCCTCA
TGCTACCAAGATCTTCGTCAACGGTGTCTGGGTGGGCATTCACCAGGACCCCAAGCA
TCTCGTCCAGCAGGTCGTGGACACTCGTCGTAAATCCTACCTGCAGTACGAGGTCTC
TCTCGTCAGAGAAATTCGAGACCAAGAGTTCAAGATCTTCTCGGATGCTGGCCGTGT
CATGCGACCCGTTTTTACCGTCCAGCAAGATGAAGAGTCGGACACTGGCATTCCAAA
GGGCCACTTGGTACTGACCAAAGACCTCGTTAATAAGTTGGCCCAAGAGCAGGCCG
AGCCGCCAGAAGACCCAAGCATGAAGATTGGATGGGAGGGGCTCATCAGGGCTGGT
GCAGTTGAGTATCTCGATGCCGAGGAAGAGGAGACGGCCATGATTTGCATGACTCC
CGAGGATCTCGAGCTGTATCGTGCCCAGAAGGCAGGCATTGCCACCGAAGAGGACG
TGGGCGACGATCCGAACAAGCGACTCAAGACGAGGACAAACCCAACAACGCACAT
GTACACGCACTGCGAGATTCATCCAAGCATGATCTTGGGTATCTGCGCGAGCATCAT
TCCTTTCCCCGATCACAACCAGTCCCCCCGTAACACCTACCAATCTGCCATGGGTAA
ACAAGCCATGGGTTTCTTCCTCACCAACTATTCCAGGCGCATGGATACCATGGCCAA
CATTCTCTACTACCCCCAGAAGCCGCTGGGTACGACCCGCTCTATGGAGTTTTTGAA

GTTCCGCGAGCTGCCGGCTGGACAAAACGCCATTGTAGCCATTGCTTGTTACTCTGG

CTACAACCAGGAAGATTCGGTCATCATGAATCAGAGTAGTATTGACAGAGGCCTCTT

CCGAAGTCTCTTCTTCCGATCCTACTCGGATCAGGAGAAGAAGGTGGGCTTGAACTA

CACCGAAGTGTTCGAGAAGCCGTTCCACCAAAACACCCTCCGTATGAAGCACGGCA

CATACGACAAGCTTGACGAGGACGGCATCGTCGCTCCTGGCGTCCGAGTTTCCGGG

GAGGACATCATCATCGGCAAGACTGCGCCGATTGACGCCGACACGCAAGATCTCGG

CACCAGAACCACAATGCACCAGAGGCGTGATATCTCGACACCCCTGCGCAGCACCG

AGAACGGCATCGTGGACTCTGTCATTGTGACGGTCAACGCGGACAATGTCAAGTAT

GTCAAGGTCCGCGTCCGCACGACCAAGATTCCGCAGATTGGTGACAAGTTTGCGTCT

CGTCACGGACAAAAGGGCACTATTGGTGTCACTTACCGGCAGGAGGACATGCCGTT

CACCAGAGAGGGAATTACGCCCGACATCATCATCAACCCCCACGCCATTCCGTCTCG

TATGACAATCGCTCACTTGATTGAGTGTCTCCTAAGCAAGGTCTCTACGTTGGAGGG

TATGGAGGGTGATGCTACCCCCTTTACCGATGTCACGGTCGACTCGGTTTCGGAGCT

GCTGCGAAAGCACGGCTACCAGTCTCGAGGCTTCGAGATCATGTACAATGGTCACA

CGGGCCGCAAGCTGAGGGCCCAGGTCTTCTTTGGACCAACGTACTACCAGCGACTCC

GCCACATGGTGGACGACAAGATCCACGCCCGTGCCCGTGGTCCGGTGCAGATCATG

ACGCGACAGCCGGTGGAGGGTCGTGCCCGAGACGGTGGTCTGCGATTCGGAGAAAT

GGAGCGTGATTGCATGATTGCACACGGAGCCGCGTCCTTCCTCAAGGAGCGTTTGTT

TGAGGTGTCTGACGCCTTCCGAGTTCACATTTGCGAGATTTGTGGACTCATGACGCC

CATTGCCAACCTCTCCAAGCAATCGTTCGAGTGCCGGCCGTGCAAGAACAAGACCA

AGATTGCGCAGATTCACATCCCTTATGCGGCCAAGCTCTTGTTCCAGGAGCTCCAGT

CGATGAACATTGCCGCCCGAATGTTTACTGACCGGTCTGGCGCGTCTGTCAGGTAA

***>Ta0020_005632-RA_rpb2***

ATGGCTGATTACGAAGACGATTACGACTATGAGAACTATGGGGATGAGGATGAGGG

CATCACGCCCGAGGATTGCTGGACTGTGATTTCCTCCTTCTTCGAGACCAAGGGCCT

CGTATCGCAGCAGACCGACTCCTTTGACGAGTTCACCCAGACGACAATTCAGGATCT

CGTCAACGAATACTCCACCATCACACTCGACCAGCCCAATCCTCCTTCGCCACCTGG

TCGAACGATAGCCCTTCGCCGATATGAAATCAAATTTGGAAGCGTCATGGTATCACG

TCCCACTATCAGTGAGACGGATGGAACTGTGACGTCTTTGCTTCCTTACGAATGCCG

AGACCGCAACCTGACTTACGCCAGTCCGCTTTACATCAAGATCACCAAGAAAGTGTC

TGCGGCCGTCGAGAGGGAGGTTCCGCTGCACGAGATGGACGATGCCCAGCAGGAAC

AGTATGCAAGGACCGGAGAAAACCCCACAAAGCTGGAATGGGAGGAGGAAGAGAA

TGGCGAGGACGACAATCTCGGCAAGTCTGACGACTGGAAGGACATGGTTTTCGTTG

GCAAGCTGCCCATCATGGTCAAGTCCAAGATTTGTCATTTGAGCCGTGAACAGGATG

ACAGCCTGTTCCTCGTCAACGAGTGCCCTTACGACCAAGGAGGCTACTTTGTTATCA

ACGGTAGTGAAAAGGTCCTCATCGCCCAGGAGCGTTCCGCCGCAAACATCGTCCAG

GTCTTCAAGAAGGCCCAGCCCAGTGCTTATACCTACACGGCCGAAATCCGAAGTGC

GCTGGAAAAGGGATCTCGACTCATCTCTAGCATGATGCTCAAGTTGTATGGCAAAGG

AGACTCTGCGCGAGGTGGCTTTGGGCAAACTATTCACACTACCCTGCCTTTTGTCAA

GTCGGATCTTCCCGTTGCCATTGTTTTCCGTGCCTTGGGCGTCGTTTCTGATGAGGAC

ATTCTGAACCACATCTGCTACGACCGAAACGACAGCCAAATGCTTGAAATGCTTCGG

CCTTGCATTGAAGAGGCCTTTTGTGTTCAGGATCGAGAAGTTGCTCTTGATTTCATCG

GAAAGCGAGGCAATCGTGATCAAGCCGGCCTCGGTCGCGAGAAGCGTGTTCGTGTA

GCAAAGGACATTCTTCAGAAGGAGACGCTTCCCCACATTTCCCAGACTGAAGGCAG

TGAGACCAGAAAGGCATTCTTCCTTGGATACATGGTGCACAAGCTATTGCAATGCGC

ACTCGGAAGACGAGAGCCCGACGACCGAGATCACTTTGGAAAGAAGCGTCTGGATC

TGGCGGGTCCACTGCTGGCCAAGCTGTTCCGTGGCATCATGCGCAGAATGAACACTG

AGCTGGCCAACTACCTGAGACGATGTGTTGAGGGTAACCGCCACTTCAATCTTGCTG

TTGGCATCAAGCCCGGCACACTTTCCAACGGACTCAAGTACTCACTCGCCACTGGAA

ACTGGGGTGACCAGAAGAAGGCAATGAGCTCGACCGCAGGTGTCTCACAGGTGCTT

AACCGTTACACCTTTGCTTCTACACTTTCCCATTTGCGTCGTACCAATACACCCATCG

GAAGAGATGGTAAGCTGGCGAAGCCTCGACAGCTCCACAACACACACTGGGGCTTG

GTGTGCCCGGCTGAGACCCCTGAAGGGCAGGCTTGTGGTCTGGTCAAGAATTTGTCT

CTGATGTGCTACGTCAGTGTTGGATCTCCCTCTGAGCCTTTGATCGAGTTTATGATCA

ACAGAGGTATGGAAGTCGTCGAGGAGTATGAGCCACTGAGGTATCCCCATGCCACA

AAGATCTTTGTGAATGGTGTCTGGGTTGGAATCCATCAAGACCCCAAGCATCTGGTA

AACCAAGTCTTGGATACTCGTCGCAAATCCTATCTGCAGTACGAAGTCTCTCTGATC

AGAGAAATCCGAGACCAAGAATTCAAAATCTTCTCTGATGCCGGTCGTGTTATGCGT

CCCGTCTTCACTGTGCAGCAGGAAGATGACCCGGAAACGGGTATCAACAAGGGCCA

CCTGGTTCTGACCAAGGACCTCGTCAATAGGCTGGCCAAAGAGCAGGCTGAGCCTC

CAGAAGACCCAAGCATGAAGCTCGGATGGGAGGGGCTGATTAGGGCTGGTGCGGTG

GAATATCTCGACGCCGAGGAAGAAGAAACATCCATGATTTGCATGACACCGGAAGA

TCTTGAGCTTTATCGTCTTCAAAAAGCCGGCATTGCCACGGATGAAGACATAGGAGA

TGACCCAAATAAGCGTCTCAAGACCAAGACAAATCCGACAACTCACATGTATACGC

ATTGCGAGATTCACCCGAGTATGATCTTAGGTATCTGTGCTAGTATCATTCCTTTCCC

CGATCACAACCAGTCCCCCCGTAACACCTACCAGTCTGCCATGGGTAAACAAGCCAT

GGGCTTCTTTTTGACCAACTATTCTCGTCGTATGGACACCATGGCCAACATCCTCTAC

TACCCTCAGAAACCGCTGGGCACCACTCGTTCTATGGAGTTTTTGAAATTCCGTGAG

CTGCCAGCCGGACAAAACGCCATTGTAGCAATTGCTTGTTACTCTGGTTATAACCAA
GAAGATTCCGTCATTATGAACCAAAGTAGTATTGACAGAGGTCTCTTCCGAAGTCTT
TTCTTCCGATCATATTCGGATCAAGAGAAGAAGGTTGGCTTGAACTACACGGAAGTG
TTTGAGAAGCCATTCCACCAAAACACTCTTCGCATGAAGCACGGCACATATGACAA
GCTGGATGAAGATGGTATCGTTGCTCCCGGTGTTCGTGTGTCAGGTGAGGATATCAT
CATCGGCAAGACAGCACCAATCGACGCCGAGACACAGGATCTTGGAACCAGGACGA
CCATGCACCAAAGACGCGATATCTCGACGCCTTTGCGAAGCACCGAGAACGGTATT
GTCGACCAGGTCATCGTGACGGTGAACGCGGATAATGTCAAATATGTCAAGGTTCGT
GTTCGAACGACTAAGATTCCCCAGATTGGAGACAAGTTTGCCTCTCGTCACGGTCAG
AAGGGCACTATTGGTGTTACCTACCGACAGGAGGATATGCCTTTCACAAGAGAAGG
TCTCACTCCAGATATCATTATCAACCCCCACGCTATTCCGTCTCGTATGACAATTGCT
CACTTGATTGAGTGTCTGCTGAGTAAGGTCTCTACGTTGGAAGGTATGGAGGGTGAT
GCTACTCCCTTTACGGATGTCACCGTCGACTCAGTCTCAGAGCTCTTGCGAAAGCAC
GGCTACCAGTCTCGGGGCTTCGAGATTATGTACAACGGCCATACTGGACGCAAGCTA
CGAGCACAAGTGTTCTTTGGACCAACATACTACCAGCGACTCCGCCACATGGTGGAC
GACAAGATCCATGCCAGAGCTCGTGGCCCTGTGCAGATCATGACACGGCAGCCAGT
GGAGGGTCGTGCTCGAGATGGTGGTCTCCGATTCGGAGAAATGGAACGTGATTGCA
TGATTGCACACGGTGCAGCGTCCTTCCTCAAGGAGCGACTGTTTGAGGTGTCGGACG
CCTTCCGAGTTCACATTTGCGAGATTTGTGGACTCATGACGCCCATTGCGAATTTATC
CAAACAATCGTTCGAGTGTCGGCCATGTAAGAACAAGACAAAGATTGCGCAGATTC
ACATTCCTTATGCTGCCAAGCTCTTATTCCAGGAGCTCCAGTCAATGAACATTGCAG
CTAGAATGTACACCAACCGGTCTGGTGCATCTGTCCGGTAG

***>Th3844_ITS***

AGGGATCATTACCGAGTTTACAACTCCCAAACCCAATGTGAACGTTACCAAACTGTT

GCCTCGGCGGGATCTCTGCCCCGGGTGCGTCGCAGCCCCGGACCAAGGCGCCCGCC

GGAGGACCAACCTAAAACTCTTATTGTATACCCCCTCGCGGGTTTTTTTATAATCTGA

GCCTTCTCGGCGCCTCTCGTAGGCGTTTCGAAAATGAATCAAAACTTTCAACAACGG

ATCTCTTGGTTCTGGCATCGATGAAGAACGCAGCGAAATGCGATAAGTAATGTGAAT

TGCAGAATTCAGTGAATCATCGAATCTTTGAACGCACATTGCGCCCGCCAGTATTCT

GGCGGGCATGCCTGTCCGAGCGTCATTTCAACCCTCGAACCCCTCCGGGGGGTCGGC

GTTGGGGATCGGCCCTCCCTTAGCGGGTGGCCGTCTCCGAAATACAGTGGCGGTCTC

GCCGCAGCCTCTCCTGCGCAGTAGTTTGCACACTCGCATCGGGAGCGCGGCGCGTCC

ACAGCCGTTAAACACCCAACTTCTGAAATGTTGACCTCGGATCAGGTAGGAATACCC

GCTGAACTTAA

***>Th0179_ITS***

AGGGATCATTACCGAGTTTACAACTCCCAAACCCAATGTGAACGTTACCAAACTGTT

GCCTCGGCGGGATCTCTGCCCCGGGTGCGTCGCAGCCCCGGACCAAGGCGCCCGCC

GGAGGACCAACCAAAACTCTTATTGTATACCCCCTCGCGGGTTTTTTTATAATCTGA

GCCTTCTCGGCGCCTCTCGTAGGCGTTTCGAAAATGAATCAAAACTTTCAACAACGG

ATCTCTTGGTTCTGGCATCGATGAAGAACGCAGCGAAATGCGATAAGTAATGTGAAT

TGCAGAATTCAGTGAATCATCGAATCTTTGAACGCACATTGCGCCCGCCAGTATTCT

GGCGGGCATGCCTGTCCGAGCGTCATTTCAACCCTCGAACCCCTCCGGGGGGTCGGC

GTTGGGGATCGGCCCTCCCTTAGCGGGGGCCGTCTCCGAAATACAGTGGCGGTCTCG

CCGCAGCCTCTCCTGCGCAGTAGTTTGCACACTCGCATCGGGAGCGCGGCGCGTCCA

CAGCCGTTAAACACCCAACTTCTGAAATGTTGACCTCGGATCAGGTAGGAATACCCG

CTGAACTTAA

>***Tr0711_ITS***

CGTTACCAATCTGTTGCCTCGGCGGGATTCTCTGCCCCGGGCGCgTcgcAgCCCCGGAT

CCCATGGCGCCCGCCGGAGGACCaaCTCAAACTCTTTTTCTCTCCGTCGCGGCTTCC

GTCGCGGCTCTGTTTTACCTTTGCTCTGAGCCTTTCTCGGCGACCCTAGCGGGCGTCT

CGAAAATGAATCAAAACTTTCAACAACGGATCTCTTGGTTCTGGCATCGATGAAGAA

CGCAGCGAAATGCGATAAGTAATGTGAATTGCAGAATTCAGTGAATCATCGAATCTT

TGAACGCACATTGCGCCCGCCAGTATTCTGGCGGGCATGCCTGTCCGAGCGTCATTT

CAACCCTCGAACCCCTCCGGGGGGTCGGCGTTGGGGATCGGCCCCTCACCGGGCCG

CCCCCGAAATACAGTGGCGGTCTCGCCGCAGCCTCTCCTGCGCAGTAGTTTGCACAC

TCGCACCGGGAGCGCGGCGCGGCCACAGCCGTAAAACACCCCAAACTCTGAAATGT

TGACCTCGGATCAGGTAGGAATACCCGCTGAACTTAAGCATATCAA

>***Ta0020_ITS***

GTGAACCATACCAAACTGTTGCCTCGGCGGGGTCACGCCCCGGGCGCGTCGCAGCC

CCGGAACCAGGCGCCCGCCGGAGGGACCAACCAAACTCTTTACTGTAGTCCCCTCG

CGGACGTTATTTCTTACAGCTCTGAGCAAAAATTCAAAATGAATCAAAACTTTCAAC

AACGGATCTCTTGGTTCTGGCATCGATGAAGAACGCAGCGAAATGCGATAAGTAAT

GTGAATTGCAGAATTCAGTGAATCATCGAATCTTTGAACGCACATTGCGCCCGCCAG

TATTCTGGCGGGCATGCCTGTCCGAGCGTCATTTCAACCCTCGAACCCCTCCGGGGG

GTCGGCGTTGGGGACCTCGGGAGCCCCTAAGACGGGATCCCGGCCCCGAAATACAG

TGGCGGTCTCGCCGCAGCCTCTCCTGCGCAGTAGTTTGCACAACTCGCACCGGGAGC

GCGGCGCGTCCACGTCCGTAAAACACCCAACTTCTGAAATGTTGACCTCGGATCAGG

TAGGAATACCCGCTGAACTTAAGCATATCA
